# A lizard is never late: squamate genomics as a recent catalyst for understanding sex chromosome and microchromosome evolution

**DOI:** 10.1101/2023.01.20.524006

**Authors:** Brendan J. Pinto, Tony Gamble, Chase H. Smith, Melissa A. Wilson

## Abstract

In 2011, the first high-quality genome assembly of a squamate reptile (lizard or snake) was published for the green anole. Dozens of genome assemblies were subsequently published over the next decade, yet these assemblies were largely inadequate for answering fundamental questions regarding genome evolution in squamates due to their lack of contiguity or annotation. As the “genomics age” was beginning to hit its stride in many organismal study systems, progress in squamates was largely stagnant following the publication of the green anole genome. In fact, *zero* high-quality (chromosome-level) squamate genomes were published between the years 2012–2017. However, since 2018, an exponential increase in high-quality genome assemblies has materialized with 24 additional high-quality genomes published for species across the squamate tree of life. As the field of squamate genomics is rapidly evolving, we provide a systematic review from an evolutionary genomics perspective. We collated a near-complete list of publicly available squamate genome assemblies from more than half-a-dozen international and third-party repositories and systematically evaluated them with regard to their overall quality, phylogenetic breadth, and usefulness for continuing to provide accurate and efficient insights into genome evolution across squamate reptiles. This review both highlights and catalogs the currently available genomic resources in squamates and their ability to address broader questions in vertebrates, specifically sex chromosome and microchromosome evolution, while addressing why squamates may have received less historical focus and has caused their progress in genomics to lag behind peer taxa.

## History and Background

Genome sequencing has revolutionized biology in every group of organisms; however, some organismal groups have better representation, genomically, than others. In the intervening years between the first lizard karyotype (Tellyesniczky, 1897) and first published lizard genome (Alföldi et al. 2011), many questions have been raised where squamate reptiles stand to provide unique insight into the patterns and processes of genome evolution including those character states shared with other organismal groups (e.g. Perry et al. 2021; Pinto et al. 2019a) and those unique to squamates (e.g. Gamble 2019). Namely, squamates provide an invaluable model system for two areas of active research: (1) the evolution of sex chromosomes (Gamble et al. 2015a) and (2) the evolution and function of microchromosomes (Perry et al. 2020). We start by briefly reviewing the development of the history of squamate genomics since its inception.

The argument for why sequencing lizard genomes is necessary, as a departure from human- and laboratory model-centric taxa, was first made in 2005 (Losos et al. 2005). Five years later, the green anole (*Anolis carolinensis*) genome appeared on NCBI and the paper published the following year (Alföldi et al, 2011). However, genomics in squamate reptiles (lizards and snakes) has lagged behind most other vertebrate groups and *all* other amniote lineages (Hotaling et al. 2021). Another seven years passed until the second high-quality squamate genome was made available through the intervention of the DNAZoo sequencing initiative, with the re-scaffolding of the Burmese python (*Python bivittatus*) genome into a chromosome-level assembly (Figure 1; Castoe et al. 2013; Dudchenko et al. 2017 & 2018). Herein, we roughly define “high-quality” genomes as those scaffolded into representative chromosomal linkage groups (scaffolds) but acknowledge that this ignores the contiguity of the primary assembly (contigs), which is possibly more important for assembly accuracy and suggest readers incorporate this metric when both assembling/publishing new assemblies or choosing an available assembly for use. As of July 12th, 2022, we had identified 73 ‘publicly available’ genome assemblies across squamate reptiles, 81% of which were published in the last 5 years (2018–present). Further, it’s been as many years since the last review of squamate genomics (Deakin and Ezaz, 2019). Due to this lag behind other vertebrate groups, such as birds—who recently surpassed 500 genome assemblies (Bravo et al. 2021) for the approximately 11,162 available bird species, squamates have largely been overlooked as a key evolutionary group for genomics studies, with ∼11,300 species until just recently represented by the lone *Anolis carolinensis* genome (Hotaling et al. 2021; Rhie et al. 2020; Uetz et al. 2022). Thus, to help refresh this mindset, we provide an up-to-date review to acclimate scientists, from taxonomically-focused biologists to computational biologists, on the state of genomics within squamate reptiles—a key, yet understudied, model group to address important biological questions in an evolutionary context.

**Figure 1:**
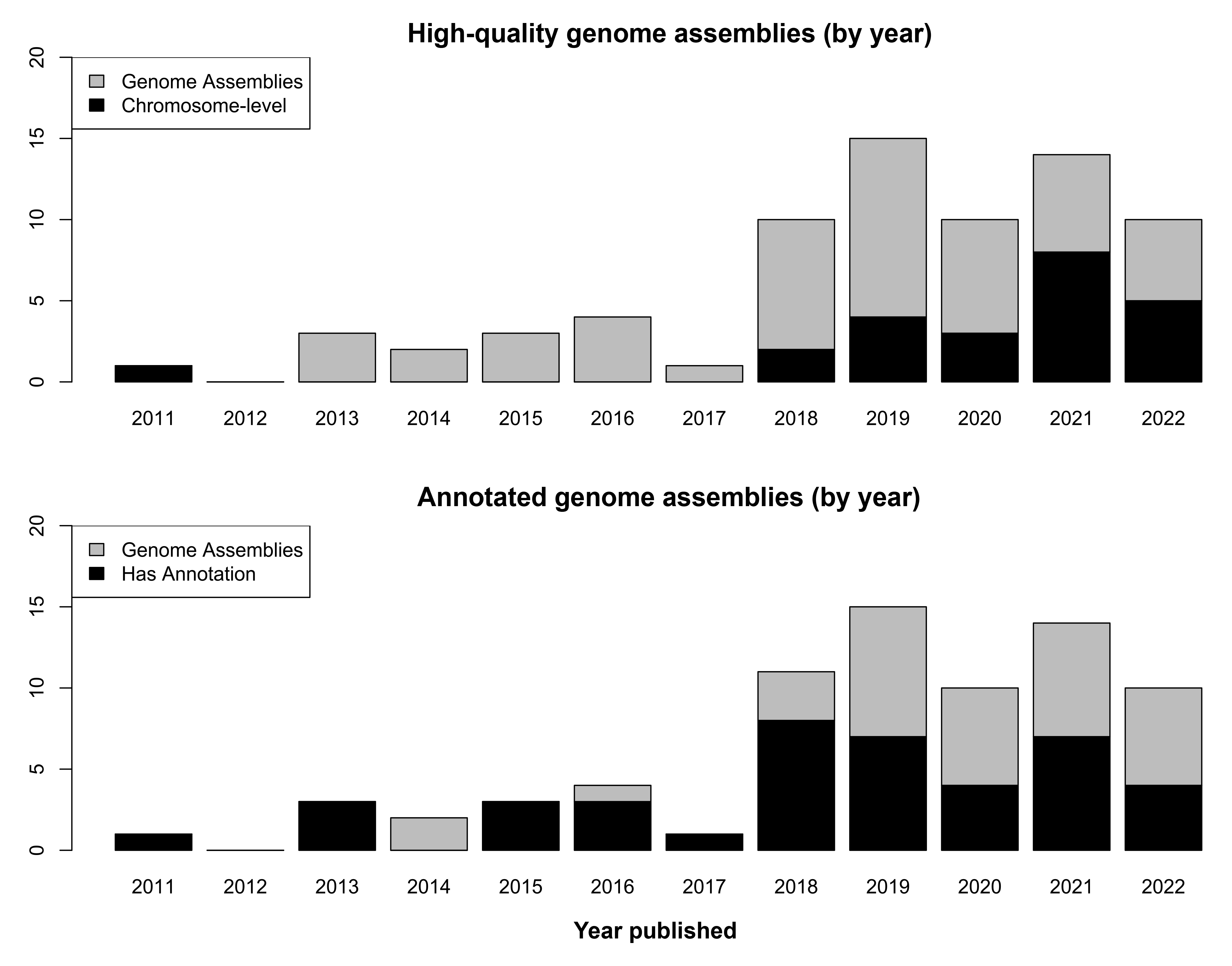
Chronological breakdown of genome assemblies published per-year and proportion of the assemblies that are chromosome-level (top panel) or annotated (bottom panel). Importantly, not all chromosome-level genomes are annotated and most chromosome-level assemblies that improve a previously annotated assembly do not publish updated annotations.

## Squamate Genomics Today

In Appendix I, we aggregated a near-complete list of squamate genome assemblies and assembly information to (1a) characterize why squamate genome assemblies have lagged behind other groups and (1b) identify specific taxonomic groups within the field that are lacking, (2) interrogate various assembly metrics across taxa to identify potential trends in data generation and assembly, and (3) discuss how currently available squamate genomes, although lacking in phylogenetic density (number of taxa), still possess the phylogenetic breadth to revise how we think about vertebrate genomics, specifically (3a) sex chromosome evolution and (3b) microchromosome evolution. As of mid-2022 (the data collection cutoff date for this manuscript), among all available squamate genome assemblies, snakes outnumbered all others combined (37 snake vs. 34 lizard assemblies). However, when accounting for only high-quality assemblies the numbers reverse (9 snake vs. 16 lizard assemblies). Importantly, all but one of these assemblies was published in the last five years (Figure 1).

One important factor in the historical lag in squamate genomics behind other amniotic groups is likely, at least in part, due to faith placed in large-scale sequencing initiatives that have then prioritized other groups. In short, the future of high-quality squamate genome generation is in the hands of those with a keen interest in reptiles. Large-scale sequencing initiatives with large resource pools, such as the Vertebrate Genome Project (VGP) consortium, have largely neglected this speciose group of amniotes (Genome 10K Community of Scientists, 2009). For scale, according to the IUCN Red List (i.e. Uetz et al. 2022), there are more non-avian reptiles (11,690) than avian (birds; 11,162)—even approaching twice as many species of squamates (11,300) than mammals (6,578)—however, as of mid-2022 of the 129 amniote genomes available through the VGP 33% (43/129) were birds and 21% (27/129) were mammals, with a staggering 1.5% (2/129) and 3% (4/129) for squamates and non-avian reptiles, respectively (https://hgdownload.soe.ucsc.edu/hubs/VGP). Without changes to these trends, there appears to be little hope for squamate genomes to be generated *en masse* through these types of initiatives. Funding agencies appear to be responding to this need and funding genome projects by smaller research groups who are excited about, and committed to, assembling reptile genomes (authors pers. obs.).

One issue that continues to inhibit accurate characterization and analysis of squamate genomes broadly, is the lack of centralization, or even a semi-centralization, of the available genomic resources (Appendix I). While most genomes have made their way to NCBI’s GenBank or other international government-sponsored analogs (e.g., ENI, CNCB), many remain scattered throughout unincorporated repositories that remain difficult to track down *a priori* (e.g., Figshare, GigaDB, DNAZoo, etc.). However, we believe this issue is larger than researchers simply not wanting to centralize these data for broader ease of access. From a researcher perspective, submitting a genome to GenBank (or similar repository) is a non-trivial task and becomes extremely cumbersome when attempting to accompany the genome assembly with annotation information generated “in-house”. Indeed, while it is a trivial task to upload a gzipped FASTA and GFF file to a third-party data repository (e.g. Figshare), or even simply a FASTA genome file to Genbank, uploading the GenBank-specific formatted assembly/annotation has multiple challenges. For example, most annotation programs don’t generate the required files for downstream use, and it then falls on researchers to then generate these files post hoc, opt for a third-party data repository to save significant time and effort, or some hybrid between the two—with the assembly cataloged on a government server and the annotation housed in a third-party repository. Although there are a few available programs that attempt to bridge this gap by piping necessary annotation software together, they are not without their difficulties (Banerjee et al. 2021; Cantarel et al. 2008; Hoff et al. 2019; Palmer, 2018). There remains a need to centralize genome assemblies with consistent, high-quality annotation information. At present, the ideal situation appears to be submitting a genome to NCBI and inquiring with RefSeq about providing annotation, which will generally provide high-quality genome annotations assuming sufficient RNAseq data is available. However, this avenue can only progress once the genome has been publicly released and can take many months due to an ever-growing queue. Accompanying high-quality genome assemblies with complimentary genome annotation is essential for drawing significant biological insights from new high-quality, reference genome assemblies. Thus, we must forewarn that although the increased quality of DNA sequencing technologies and genome assembly tools have caused a ‘boom’ in genome assembly generation across the tree of life, the subfield of squamate genomics may ‘bust’ under its own weight if steps are not taken soon to address the laborious nature of genome annotation and data dissemination. We see potential avenues for cloud computing to lessen this burden for individual research groups as databases, such as NCBI’s Sequence Read Archive (SRA), move to becoming available on the cloud (https://anvilproject.org/ncpi). It is widely known that NCBI’s GenBank, for example, provides extensive curation services and continues to expand its functions and utility, such as recently adding the NCBI Datasets (https://www.ncbi.nlm.nih.gov/datasets/) for querying data across studies and the Comparative Genome Viewer (https://ncbi.nlm.nih.gov/genome/cgv) for understanding synteny across reference assemblies. We hypothesize that these functions will only increase in utility if the activation energy for data uploading were to be reduced in some way.

## (1a) Why have squamate genomics lagged behind other groups?

Two major factors appear to have synchronously contributed to the lag in squamate genome sequencing relative to other vertebrate groups: genome size and funding. For most vertebrate groups, genomic investigations have benefited from either small genome sizes (i.e. could accomplish more with less) and/or substantial funding models (i.e. could accomplish more with more). For example, birds (637 assemblies representing 11,162 species; Bravo et al. 2021) and fishes (594 assemblies representing 32,000 species; Randhawa & Pawar, 2021) each possess some of the smallest vertebrate genomes described—most within the ∼0.4-1.4Gb range. While far larger genome sizes occur in mammals (∼2.5-3.5Gb), applied funding from health and agricultural sources (far exceeding that allocated to other vertebrate groups, such as squamates) have offset similar phenomena in the field of mammal genome sequencing (Supplemental Table 1). In the most extreme case, amphibian genomes are even larger and suffer more greatly than squamates due to this form of genome size bias, however, further extrapolation here is beyond the scope of the current article. At a glance, squamates have an average genome size of 1.73Gb (*N*=71), ranging from 1.1Gb in *Crotalus pyrrhus* assembly (Gilbert et al. 2014) to 2.86Gb in *Sceloporus occidentalis*. However, this estimate is fraught with bias due to an overabundance of low-quality short-read assemblies that likely skew the genome size estimates lower than reality (Supplemental Figures 2 and 3). We can roughly account for this by discarding all genome size estimates from primary assemblies derived from short-read technologies, assuming long-read primary assemblies are better representations of the repeat content within a genome (Rhoads and Au, 2015). This provides a revised estimate of the approximate average genome size in squamates of 1.86Gb (*N*=17), ranging from 1.39Gb in *Lacerta agilis* to 2.86Gb in *S. occidentalis*. Thus, the larger genome sizes in squamates, albeit on average still ∼0.8-1Gb smaller than mammals, combined with less overall funding than mammalian taxa, has likely led to a stagnation in high-quality genome assemblies in squamates—that is until the cost of sequencing decreased exponentially over the past five years (Wetterstrand, 2021). Thus, as sequencing costs have declined exponentially, requiring less funding to accomplish more sequencing, the subfield of squamate genomics has finally erupted and is beginning to flourish (Figure 1).

**Figure 2:**
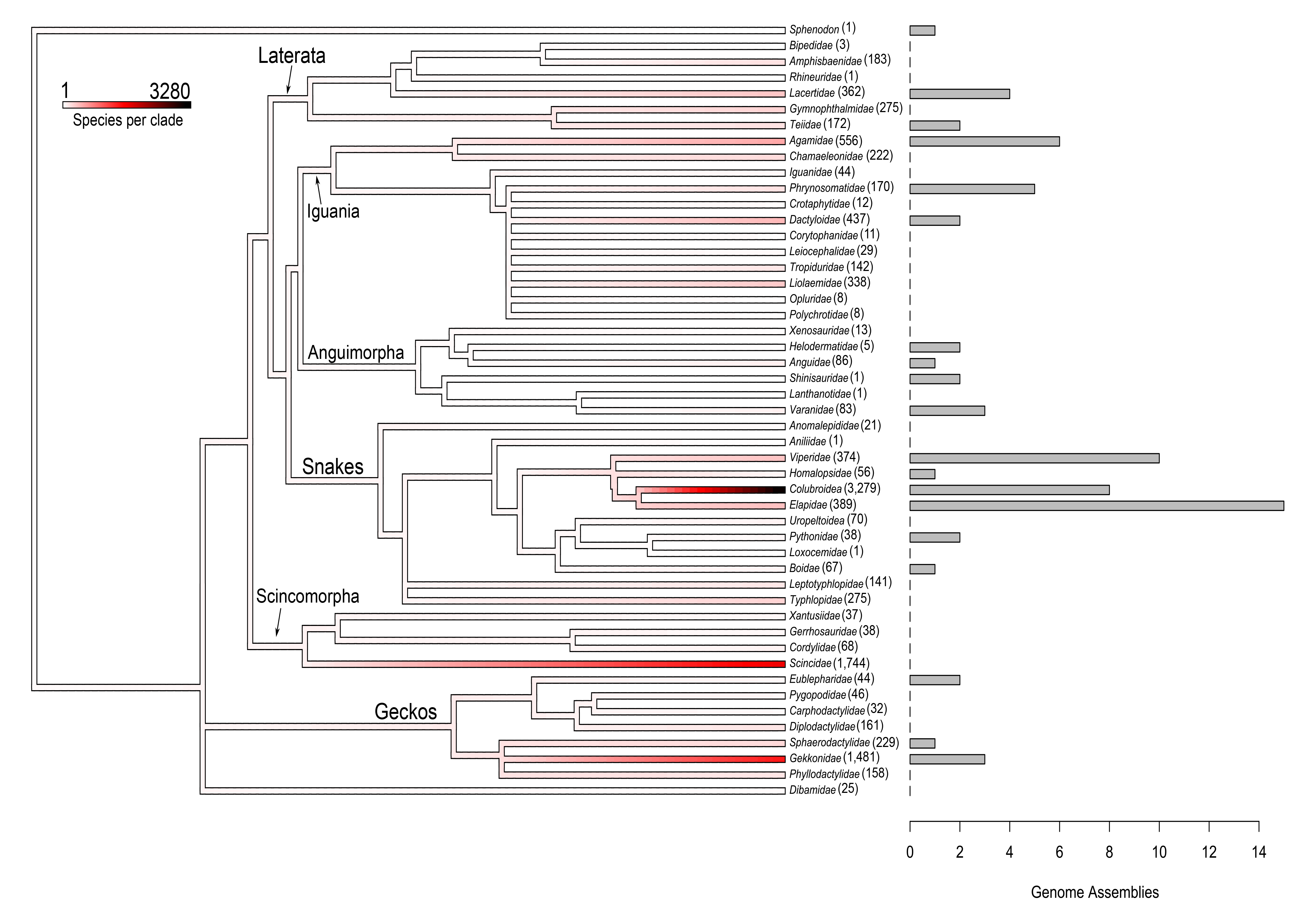
Breakdown of total number of published genome assemblies (bar graph) per phylogenetic group (family or superfamily) co-plotted with number of species in each clade (branch colors and parenthetical numbers). Phylogeny from TimeTree using a representative species from each clade (Kumar et al. 2017), species counts from the Reptile Database (Uetz et al. 2022), and plotted using phytools (Revell, 2012).

**Figure 3:**
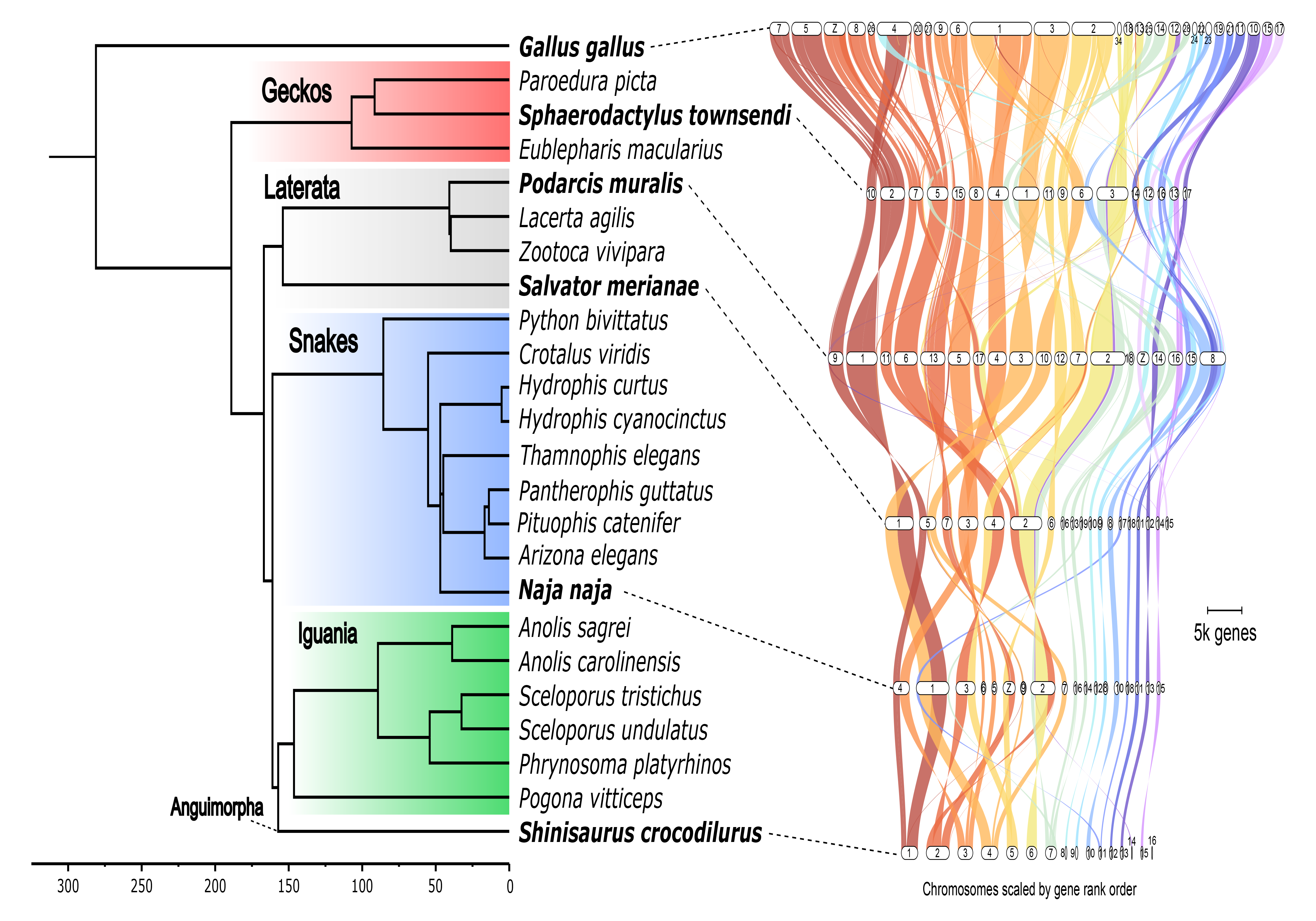
Time-calibrated phylogeny of squamate reptiles pruned to include only species with high-quality genome assemblies (rooted with chicken, *Gallus gallus*). Branches leading to major phylogenetic groups labeled, those with multiple taxa are highlighted. Phylogeny obtained from TimeTree (Kumar et al. 2017) and plotted using GeneSpace (Lovell et al. 2022) and FigTree [v1.4.4]. It’s apparent that microchromosomes are homologous in squamates that possess them (*Salvator*, *Naja*, and *Shinisaurus*), while different linkage group fusions have led to their loss in taxa that lack them (*Sphaerodactylus* and *Podarcis*).

## (1b) What taxonomic groups remain unsampled?

An overarching theme of the current state of squamate genomics is that, while few groups are adequately represented in terms of genomic resources (such as elapid snakes – 15 genomes from 395 species), most squamate groups are in dire need of additional high-quality genomic resources (such as geckos, from our very biased point-of-view, which include 2,186 species, but only six genomes). However, there are many extremely diverse and evolutionarily important groups that are completely absent, such as chameleons (222 species), amphisbaenids (182 species), and scincomorphs (1,886 species) (Figure 2). In fact, approximately five years ago, a high-quality multi-tissue transcriptome was published for the veiled chameleon (*Chamaeleo calyptratus*) with an accompanying call for additional genomic resources to be generated for this extremely interesting clade (Pinto et al. 2019b). However, to date, there has yet to be a single genome assembly of any quality, made publicly available for a chameleon—with a similar situation at play in scincomorphs and many other squamate families (Figure 2). Indeed, of the 46 squamate families that appear in Figure 2, 31 families including all chameleons and scincomorphs—occurring globally, except Antarctica—have no publicly available reference genomes. Future directions in squamate genomics should focus on including these missing taxa as important players in the investigations in the genomics of vertebrates.

## (2) Trends in data generation and assembly

Regardless of sequencing methodology implemented, most empiricists have become aware that the quality of sample collection and preparation can “make or break” a genome assembly experiment. This includes every stage of sample preparation up to its conversion from bases to bytes, including, but not limited to: tissue selection, dissection, storage, extraction, library prep, sequencing, and assembly (Dahn et al. 2022; Pinto et al. 2022 & 2023). Many squamate species are rare/hard to collect, have a limited distribution (Meiri et al. 2018), and lack material in museum collections adequate for long-read sequencing or chromatic-contact sequencing (HiC). Most species will need new specimens to be collected specifically for a genome sequencing project. Fortunately, relative to some other animal groups (for example, freshwater bivalves (Smith, 2021) and *Xiphophorus* fishes (author’s pers. obs.)), squamate DNA appears to remain remarkably stable throughout this process, which provides some relief for field collection and sub-optimal tissue conditions—preferring blood or liver tissue when available (Dahn et al. 2022; Pinto et al. 2022 & 2023). These factors prime squamates to benefit from recent advances in sequencing technology – like long, accurate sequencing reads – that have opened new doors in genome assembly. Just as the publication of the first human reference genome at the turn of the century signaled the beginnings of the “genomics age”, the recent publication of the complete human telomere-to-telomere (T2T) genome assembly has signaled a ‘rebirth’ of the genomics age, where now all model systems can be subject to high-quality reference genomes for relatively low cost, including squamates (Nurk et al. 2022; Pinto et al. 2022, 2023; Rhie et al. 2021; Sun et al. 2021). Recent advances in increasing contiguity of primary genome assemblies has been driven by third generation sequencing technologies, including Pacific Biosciences (PacBio) and Oxford Nanopore platforms (e.g. Nurk et al. 2022; Peona et al. 2020). For the past two years, PacBio High Fidelity (HiFi) reads have shown that high-accuracy reads (∼20 kb; phred quality scores ∼20+) can outperform longer reads with lower accuracy (∼40 kb+; phred quality scores ∼10) in many cases— certainly at the cost-per-base (Lang et al. 2020; Peona et al. 2020; Vollger et al. 2020). However, recent data also confirmed that some genomic regions require ultra-long read lengths to overcome extremely long stretches of repetitive DNA, some lengths of which may still be unachievable, but certainly enforces a hard ceiling for the ‘coverage-to-contiguity’ ratio at around 30X when using HiFi data alone (Pinto et al. 2023; Sun et al 2021). That said, Oxford Nanopore’s forthcoming Q20 chemistry (Kit 14 with the R10.4.1 flowcell) may provide the missing link in completing Telomere-to-Telomere (T2T) genome assemblies that makes them more approachable to squamate researchers on a tight budget.

One way of accelerating genome assembly generation across squamates would be to decentralize sequencing and assembly. This is currently how squamate genomics has advanced and assisting the field in this endeavor is one goal of this manuscript. However, it is far from as decentralized as one might imagine. Indeed, when delving into where lepidosaur (squamates and the tuatara) genome assemblies are derived from, the research group doing the sequencing (inferred via first and last authorships), and the vast majority of assemblies come from research groups in the ‘global north’ (75%; 55/73) and China (21%; 15/73); totaling 96% of all assemblies (70/73). This leaves only three assemblies having been generated by the rest of the global community. Importantly, these numbers *do not* account for the middle-author contributions to projects made by members of the global south, which are no doubt significant. In response to these types of devastating numbers, organizations, such as GetGenome (getgenome.net), may be helpful in reducing this disparity between the global north and south. Importantly, organizations like this that formally empower groups to conduct this work in-house, instead of outsourcing to a consortium, will likely produce greater innovation using these data in the long run (e.g. Hofstra et al. 2020) and could help more broadly mitigate the current state of scientific exclusion of the global south within the subfield. As such, it is important to note that since the subfield of squamate genomics is relatively young we are in an optimal position to lead an equitable globalization effort moving forward with regard to data generation and usage—an important steppingstone for the herpetological field more broadly.

### Practical considerations regarding sex chromosomes in squamate genomics

More generally, as genome sequencing technologies are capable of producing both long and accurate sequence reads, an important step to genome assembly is producing fully phased, or haplotype-resolved, genome assemblies in place of traditional chimeric assembly where alleles are assembled together (Cheng et al. 2021 & 2022). This may allow for the resolution of divergent genomic regions of biological importance, such as polyploid genomes, heterozygous inversions, alternative splice variants, and sex chromosomes (XY or ZW). Indeed, once haplotype-resolved genomes become common within squamates, sex chromosomes within the assemblies will be phased—as they are in the human genome and some others (Nurk et al. 2022; Webster et al. 2019).

Once both sex chromosomes (X and Y or Z and W) are present in the reference assembly, researchers will need to specifically assess and account for the sex chromosome complement when conducting bioinformatic experiments, such as read mapping and variant calling (Carey et al. 2022; Olney et al. 2020; Pinto et al. 2023b; Webster et al. 2019). To effectively account for sex chromosome complement in an assembly, haplotypes of the sex chromosomes must be resolved. In tandem with this paper, the first attempt at generating a haplotype-resolved genome of a squamate, a temperature-dependent sex determining gecko—the leopard gecko, *Eublepharis macularius*, was published (Pinto et al. 2023). With this along with other recent assemblies, the advent of reference quality, phased genomes for squamate taxa has become achievable for the average research group and bodes well for the future study of sex chromosomes across squamates.

## (3a) Genomics and sex chromosomes in squamates

Squamates are an invaluable model system for studying sex chromosome evolution. Within their ranks all three major modes of vertebrate sex determination occur: environmentally determined sex (temperature dependence) and genetic sex determination (both male heterogamety [XX/XY] and female heterogamety [ZZ/ZW] systems), with multiple independent transitions among the three mechanisms (Gamble et al. 2015a; Stöck et al. 2021). Studying squamates provides a powerful system to better understand the gaps in our knowledge of sex chromosome evolution broadly; specifically questions such as, (I) are some linkage groups more likely to be recruited as a sex determining role than others?, (II) are ancient sex chromosome systems an evolutionary trap that species cannot escape?, and (III) how do mechanisms of dosage balance and compensation between the sexes evolve? (e.g. Gamble et al. 2015a; Kratochvíl et al. 2021; Nielsen et al. 2019; Rupp et al. 2017). We explore these topics framed by how modern squamate genomics stand to help answer these questions.

### (I) Are some linkage groups more likely to be recruited as a sex determining role than others?

The identification and characterization of sex chromosome systems are perhaps the most well-reviewed aspect of squamate genomics—whose study has also been intimately associated with the advent of genomics in squamates—with progress increasing exponentially in recent years (Gamble, 2010; Gamble et al. 2015a, 2017, 2018; Kratochvíl et al. 2021; Pinto et al. 2022; Stöck et al. 2021). Four species-rich clades, with known, conserved sex chromosome systems – pleurodonts (iguanas, spiny lizards, and anoles, excluding corytophanids); caenophidian snakes; skinks; and lacertids – make up approximately 60% of squamate species (Rovatsos et al. 2014, 2015, 2019a; Nielsen et al. 2019; Kostmann et al. 2021). The remaining 40% of squamate species are in clades with varying levels of sex chromosome conservation, although transitions are likely common in many of these groups (Gamble et al. 2015a; Gamble et al. 2017; Nielsen et al. 2018; Keating et al. 2022; Pinto et al. 2022). Given the available data it has been suggested that linkage group recruitment as sex chromosomes is nonrandom, i.e. some linkage groups are more likely to be recruited as a sex chromosome than others (Kratochvíl et al. 2021). However, the pattern was weak and the discovery of additional linkage groups acting as sex chromosomes in geckos and dibamids requires a re-evaluation (Pensabene et al. 2023; Pinto et al. 2022; Rovatsos et al. 2022). Additionally, inferences that all taxa within a clade share an ancestral sex chromosome, i.e. knowing 5% of taxa sex chromosome systems and inferring that we know ∼60%, is drawn from Occam’s razor using sparse sampling (Kostmann et al. 2021), but in squamate sex chromosome evolution, where sex chromosome turnovers are commonplace, this kind of assumption has been shown to be untrue (e.g. Gamble et al. 2017). Thus, it stands to reason that this 60% figure may be an overestimate. Fortunately, recent advances in DNA sequencing technologies have allowed us to sample more broadly and ask finer-scale questions about how sex chromosomes originate, degenerate, and turnover (e.g. Acosta et al. 2019; Gamble et al. 2015a; 2017; 2018; Keating et al. 2020; Kostmann et al. 2021; Nielsen et al. 2018; 2019; 2020; Pinto et al. 2022; Rovatsos et al. 2019b; 2022); so the intertwined nature of developing squamate genomics and sex chromosome evolution presents great promise for future work in identifying and characterizing sex chromosome linkage groups across squamates.

The most conclusive evidence of shared ancestry of a sex chromosome system is the identification of a conserved primary sex determiner (or primary sex determining gene; PSD) among focal taxa (such as *Sry* in therian mammals; Graves, 2008). However, no prior publication has yet assembled the X/Z chromosome and then identified a putative PSD in a squamate, until now. Indeed, high-quality genome assemblies and annotations are only recently allowing us to confidently implicate putative PSDs in squamates. The first example to our knowledge being the Puerto Rican leaf-litter gecko, *Sphaerodactylus townsendi* (Box 1). It’s worth noting that previous implications of PSDs in squamates (*Pogona vitticeps* (*Sr1*) and anguimorphs (*Amh*): *Varanus komodoensis* and *Heloderma suspectum*) were based on incomplete catalogs of Z-linked genes (Deakin et al, 2016; Rovatsos et al. 2019; Webster et al. 2023). In an ideal world, assembling both the complete X/Z and Y/W would yield the best possible candidate PSD. Beyond implicating a candidate PSD, one downstream issue that is in the process of being overcome is that even upon the identification/confirmation of a putative PSD, we have limited capability to perform functional tests to confirm a putative sex-determining gene. Although there is significant progress happening on this front with the first successful gene editing in an *Anolis* lizard and a gecko (Rasys et al. 2019; Abe et al. 2023). High-quality genome assemblies and annotations are crucial to expanding the utility of functional genomic tools in squamates. Thus, although high-quality genomes are now allowing us to better characterize putative PSDs in squamates, we’re still a few years away from using gene editing to confirm these putative PSDs in different squamate species.

#### Box 1

##### Sex chromosomes and sex determination in squamates, a case for high-quality genome annotations

In 2021, the chromosome-level genome of *Sphaerodactylus townsendi* helped elucidate the dynamic evolution of sex chromosomes within this genus of geckos (Pinto et al. 2022). However, when examining the annotated gene content within the identified sex determining region (SDR) in the initial annotation [MPM_Stown_v2.2], we found no sign of a putative sex determining gene (gene known to have a consequential role in the vertebrate sex determining pathway). Through collaboration with NCBI RefSeq, this genome was re-annotated using only existing RNAseq data (i.e. no new transcriptomic data was generated between annotations) using the NCBI Eukaryotic Genome Annotation Pipeline. NCBI Annotation Release 100 of MPM_Stown_v2.3 provided significant improvements to the annotation quality (BUSCO completeness of annotated peptides from 61.5% to 92.5% using BUSCO [v5.1.2]). When re-examining the SDR of *S. townsendi* using this new annotation, a candidate primary sex determining gene (PSD) became clear, anti-Müllerian hormone receptor 2 (*AMHR2*). Indeed, *AMHR2* has been identified as the independently-evolved primary sex determining gene in at least two groups of fish, fugu and ayu (Kamiya et al. 2012; Nakamoto et al. 2021) and its inactivation causes male-to-female sex reversal in the Northern Pike (Pan et al. 2022). This example supports that—even without generation of additional data—high-quality annotations can be generated for divergent species with minimal available transcriptomic data. However, it is also likely that this high-level of quality for genome annotation is beyond the reach of many (if not most) biology-focused research groups, as it was for us (Pinto et al. 2022). We suggest that this may serve as motivation for the generation of the development of additional genome annotation pipelines or the adaptation of existing pipelines to be more approachable ‘lay-empiricists’ interested in answering these fundamental types of questions in newer model systems.

### (II) Are ancient sex chromosome systems an evolutionary trap that species cannot escape?

Early sex chromosome work highlighting mammalian and avian taxa suggested that perhaps ancient sex chromosome systems may become so entangled in the biology of the organisms’ development that it served as an “evolutionary trap”, to which there was little chance of escape (Bull, 1983; Bull and Charnov, 1985; Gamble et al. 2015a; Nielsen et al. 2019; Pokorná and Kratochvíl, 2009). Indeed, many taxa that possess an ancient, ancestral sex chromosome system appear to remain evolutionarily ensnared within it, including mammals (XY), birds (ZW), *Drosophila* (XY), lepidopterans (ZW), and “advanced” snakes (ZW) (Bachtrog et al. 2014; Rovatsos et al. 2015; Gamble et al. 2017; Graves, 2008; Ohno, 1967; Webster et al. 2023). To our knowledge, there are few empirical examples of taxa escaping old, degenerated sex chromosome systems (Rovatsos et al. 2019c; Terao et al. 2022). However, one possible example within squamates are the basilisk and casque-headed lizards (Corytophanidae) that possess a different sex chromosome system than all other pleurodonts. Phylogenetic uncertainty plagues this claim as a conclusive case of escaping the trap and more work is needed (Acosta et al. 2019; Nielsen et al. 2019). However, as more and more transitions among sex-determining systems have been identified it is unclear whether all sex chromosomes are destined to become traps. Because they have a variety of sex-determining systems with numerous transitions among them (Gamble et al. 2015a; Gamble et al. 2017; Pokorná and Kratochvíl 2009; Ezaz et al. 2009) squamates are an excellent model to investigate this question.

### (III) How do mechanisms of dosage balance and compensation between the sexes evolve?

Perhaps the scarcest data available regarding sex chromosomes in squamates lies in how these animals deal with gene dosage changes that evolve in response to the degeneration of the sex-limited sex chromosome. In many well-characterized animal model systems, such as the XY systems of mammals and fruit flies or the ZW systems in birds and moths, differences in gene copy number between the sex chromosomes can result in myriad disparate outcomes (Bachtrog et al. 2014; Gu and Walters, 2017; Vicoso and Bachtrog, 2009). Sex chromosome dosage work contains two interrelated questions specific to genes within the non-recombining region of the sex chromosomes, (1) what is the gene dosage between the sexes, relative to each other, known as dosage balance, and (2) what is the gene dosage of the sex chromosomes in each sex relative to the ancestral (autosomal) condition, known as dosage compensation (Gu and Walters, 2017). For instance, in mammals and moths there are mechanism(s) to globally silence one of the two X/Z chromosomes in homogametic individuals to balance the dosage between the sexes; however, although global expression between the sexes is equal, expression in both sexes is lower than the ancestral condition. In other words, mammals and moths possess dosage balance mechanisms, but not those for dosage compensation. Meanwhile, sex chromosomes in fruit flies are both balanced and compensated for, and birds are neither balanced or compensated (Gu and Walters, 2017). Because the outcomes of changes in gene dosage are disparate across taxa, more naturally occurring ‘evolutionary experiments’ are desperately needed to better understand the underpinnings of these phenomena.

Due to the lability of sex chromosomes across squamates, they may again play a pivotal role in deciphering the broader mechanistic underpinnings of sex chromosome gene dosage. Indeed, a unique characteristic of squamates relative to most other amniotes is that, due to the high rates of sex chromosome turnover, one can more easily infer the ancestral, autosomal gene expression level of multiple sex chromosome systems using closely related species (e.g. Keating, 2022) instead of using distant proxies, which may introduce additional uncertainty (e.g. Webster et al. 2023). This concept has been used across taxonomic groups to elucidate the evolutionary history of a variety of traits (e.g. Blount et al. 2018; Sackton and Clark 2019; Smith et al. 2020; Neemuchwala et al. 2023). In squamates specifically, this concept has been used to study many other independently-evolved traits such as adhesive digits (Gamble et al. 2012) and photic activity patterns (Gamble et al. 2015b; Pinto et al. 2019c), among many others. Thus, to more effectively study how dosage compensation mechanisms evolve in amniotes, squamates are an important model system to utilize. However, to-date the lack of genomic resources have especially hindered these investigations. This is a direct result of the lack of high-quality genomic resources available for squamates prior to 2018—because knowing how genes are linked together is necessary information to investigate dosage (Keating et al. 2022; Vicoso et al. 2013; Webster et al. 2023).

As stated in the introduction, the only high-quality genome available prior to 2018 was the *Anolis carolinensis* genome (Alföldi et al. 2011), as such, we know that *Anolis carolinensis* possesses both dosage balance and compensation (Marin et al. 2017; Rupp et al. 2017). Clever application of the *Anolis* genome to similar analyses in snakes also identified relatively early on that caenophidian, so-called “advanced”, snakes, like birds, lack both dosage balance and compensation (Vicoso et al. 2013), which was later confirmed using additional high-quality snake resources (Schield et al. 2019). More recently, conceptually similar approaches to those used by Vicoso et al. (2013) have led to an increase in transcriptomic data mapped to a distant relative genome to elucidate presence/absence of dosage balance in corytophanid (Pleurodonta), pygopodid (Gekkota), and anguimorph lizards (Nielsen et al. 2019; Rovatsos et al. 2019b; 2021). Additional work, including additional genomic data, have led to additional findings that both anguimorphs and diplodactylids (Gekkota), similarly to birds and snakes, appear to lack both dosage balance and compensation (Keating, 2022; Webster et al. 2023). Indeed, given the sheer diversity within Squamata our knowledge of how these animals handle dosage differences between the sexes is exceptionally sparse.

Trending with previous sections regarding the necessity of high-quality annotations to accompany high-quality genome assemblies (e.g. Box 1) also apply *ad infinitum* to studying sex chromosome dosage. When examining dosage there are essentially two scales one can use to examine differences between the sexes (a) global and (b) positional scales. (a) Global can only be used to study dosage balance at a broad scale, where comparing gene expression differences between males and females on different linkage groups (e.g. Nielsen et al. 2019; Rovatsos et al. 2019b; 2021). However, as the name might imply, this scale provides little insight into the fine-scale processes of sex chromosome evolution. Indeed, when a high-quality reference is available for a given species (or close relative) one can conduct (b) finer-scale, positional examinations of gene expression across the pseudo-autosomal (PAR) boundary and decrease noise from ‘misplaced’ genes that are no longer linked in the focal taxon, even if they are in a distant relative such as chicken or *Anolis* (Schield et al. 2019; Webster et al. 2023). Further, when examining expression on smaller chromosomes with relatively few genes, missing genes due to poor annotation quality can decrease statistical power to detect changes in dosage significantly (e.g. Keating, 2022; Webster et al. 2023). Thus in addition to addressing broader questions, such as those discussed above, high-quality annotations are necessary to accompany new reference genomes being generated to better understand how sex chromosome dosage evolves, identify putative sex determining genes (Box 1), and more generally to better characterize the “sexomes” of squamate reptiles (Stöck et al. 2021).

## (3b) Microchromosome evolution

In chicken, early 20^th^ century cytologists identified 12 easily-distinguishable large chromosomes and an additional 18+ smaller, dot-like chromosomes; Dr. Nettie Stevens notably prefaced this finding in her laboratory notebook with, “impossible to tell how many small ones” (Boring, 1923; Hance, 1924). Later work coined the term “microchromosomes” to describe these ‘innumerable’ small chromosomes and their larger counterparts as “macrochromosomes” (Newcomer, 1957; Ohno, 1961; Yamashina, 1944). However, no universally agreed upon definition of a microchromosome has yet to be established in the literature, certainly not since the advent of high-quality genome assemblies in reptiles (Boring, 1923; Fillon, 1998; Newcomer, 1957; Ohno, 1961). Indeed, at the advent of genome sequencing in birds, chicken chromosomes were arbitrarily grouped as macrochromosomes (1-5), intermediate chromosomes (5-10), and microchromosomes (11+) (Hillier et al. 2004). Subsequent studies have either used these criteria, grouping macrochromosomes and intermediate chromosomes as macrochromosomes, ranging in size from ∼23Mb to ∼200Mb (O’Connor et al. 2018), or established their own criteria for an arbitrary cutoff, such as 10Mb, 30Mb, or 50Mb (Karawita et al. 2022; Perry et al. 2021; Srikulnath et al. 2021; Waters et al. 2021). However, these arbitrary categorizations—enforced across vertebrates—make direct comparisons between taxa difficult and may encourage spurious correlations from these artifacts. These factors, among others, warrant a re-analysis of “what is a microchromosome?” and “why are they important?” and we demonstrate how squamate genomics provides vital insight into these questions.

Microchromosomes, no matter how they are defined, are present in most vertebrate groups (Srikulnath et al. 2021). However, their evolution remains murky – they have either been inherited from a common ancestor and lost independently multiple times or gained and lost independently multiple times. Since microchromosomes have historically been inhibitively difficult to assemble prior to long-read sequencing technologies, studies detailing finer-scale analyses have been lacking. Importantly, studies have lacked proper controls in an evolutionary context. No analysis to-date of microchromosomes using genomic sequence data has included, and specifically accounted for, the two squamate lineages that are known to not possess microchromosomes, i.e. geckos and lacertids (Deakin & Ezaz, 2019; Olmo, 1986; Olmo et al. 1990; Pinto et al. 2022; Srikulnath 2013; Tellyesniczky, 1897). Indeed, past studies excluding these groups have shown that microchromosomes have a set of distinct properties relative to macrochromosomes, including higher GC content, higher gene density, and a distinct nuclear architecture (Perry et al. 2021; Srikulnath et al. 2021). Here, we take a fresh look across vertebrates (mostly reptiles) as a primer to better understand the biology of microchromosomes and their evolution.

### Are microchromosomes conserved across reptiles?

Microchromosomes were likely present in the ancestor of all reptiles, including birds (Waters et al. 2021). However, within squamates, the hypothesis that the MRCA possessed microchromosomes has never been explicitly examined with synteny analyses including both geckos and lacertids. Support for an ancestral lack of microchromosomes in squamates would appear as strong conservation of linkage groups between geckos and lacertids regarding microchromosome fusions, which we do not see (Figure 3). Instead, we observe lineage-specific fusions of microchromosomes to different macrochromosomes in geckos and lacertids. Furthermore, there is a near 1:1 relationship of microchromosomal synteny across snakes, teiids, and anguimorphs—spanning the phylogenetic breadth of non-gekkotan squamates and extending to birds (Figure 3). Thus, geckos and lacertids have most likely lost microchromosomes twice independently. Additionally, when losing microchromosomes in both taxa it is apparent that, although their absolute size tends to fluctuate between taxa, their relative sizes tend to stay the same (i.e. small chromosomes tend to stay small)—unless they become fused to other chromosomes, which contrasts the patterns seen in some birds, such as chicken—which has gained multiple microchromosomes relative to the inferred ancestral karyotype (O’Connor et al. 2018). Given the currently available data, these additions to the microchromosome evolution discourse provide some insight into the evolutionary processes involved in the gains/losses of microchromosomes in certain vertebrate lineages.

### What is a microchromosome?

A null prediction of genomic composition of a chromosome might suggest, since the majority of an animal’s DNA is noncoding—all else being equal—that smaller chromosomes should have higher gene density. Similarly, GC-biased gene conversion may also lead to overall higher GC content on smaller chromosomes (Fullerton et al. 2001)—since smaller chromosomes also have less space to recombine this GC bias should, in-turn, scale with chromosome size. Therefore, to truly deviate from this null expectation, a “microchromosome” should deviate from what’s observed from closely related species that don’t possess microchromosomes. These expectations are supported by a strong linear relationship between chromosome size and gene content/GC content in species without microchromosomes, which is exactly what we see in the gecko (Pearson’s r: gene content = 0.865***/GC content = −0.781***), lacertid (gene = 0.699**/GC = −0.727**), alligator (gene = 0.936***/ GC = −0.720*), and even human (gene = 0.857***/GC = −0.598*) (Supplemental Figure 1, panels A–D).

With a null expectation between chromosome size, gene and GC content established, we examine deeper when/if deviations occur in taxa that possess microchromosomes. We find that non-avian reptiles do not deviate from the expectation of GC content based solely on chromosome size. Evidence for this observation is two-fold, (1) the overall range of GC content remains constant in non-avian reptiles, from about 42-52% genome-wide, and (2) the correlation between GC content chromosome size remains constant (Pearson’s r >-0.6) and significant in all taxa except the snake (Supplemental Figure 1, panels E–H)., which has distinct distribution of data— compared to all other taxa—with an apparent break between chromosomal GC content between 40-42% (Supplemental Figure 1, panel G). Since the extremely high GC content and presence of immensely small microchromosomes (<10Mb) in birds are both independently derived since their divergence with their closest extant relatives (crocodilians and testudines, respectively), it’s difficult to draw broader conclusions from analyzing bird genomes alone. In this context, birds also do not appear to deviate from the expectation set by other vertebrates, however, the shear diminutiveness of their microchromosomes appears to have caused them to increase GC content much higher than non-avian vertebrates have attained, ranging from 40-63% (Supplemental Figure 1, panels I–L). For birds, this excessive GC content in microchromosomes may be related to the presence of both endothermy and microchromosomes, as higher GC content is associated with thermostability of DNA molecules (Bernardi and Bernardi, 1986).

Since the advent of HiC in squamates (2018–present, e.g. Pinto et al. 2022; Shield et al. 2019) new understandings of microchromosome evolution have begun to emerge, yet are still being explored at a fundamental level. As the lines between macro- and micro-chromosomes somewhat blur in the age of chromosome-level genome assemblies, recent work has begun exploring the nuclear organization of microchromosomes in reptiles (Perry et al. 2021). Specifically, HiC data implicates a distinct intra-cellular compartmentalization of microchromosomes in the nucleus (Perry et al. 2021; Waters et al. 2021). Importantly, previous investigations were either missing data from geckos and lacertids or used arbitrary cutoffs to infer the presence of microchromosomes when they weren’t present (Figure 4).

**Figure 4:**
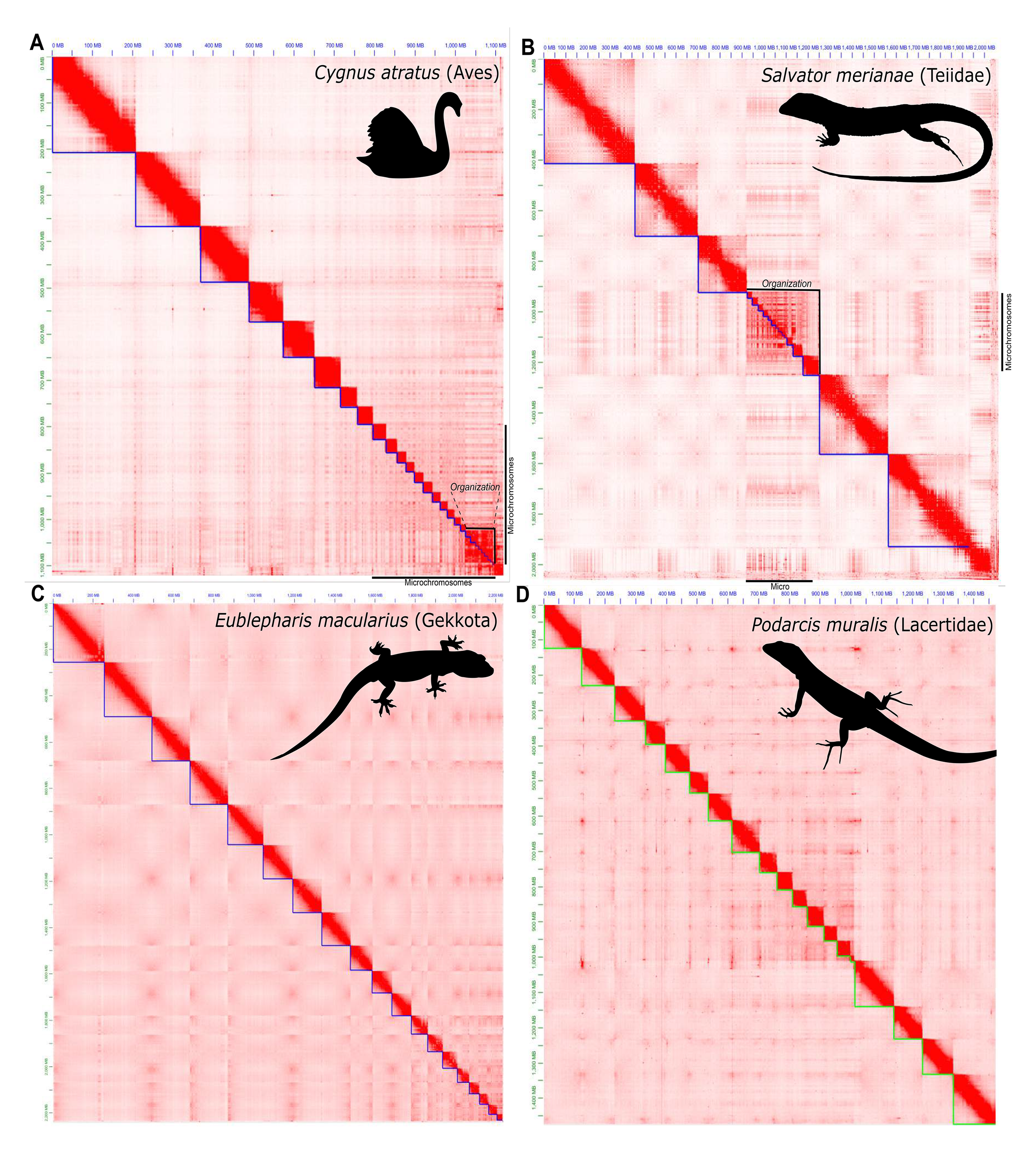
HiC contact maps for representative reptile taxa demonstrating the presence of microchromosomes in (A) birds and (B) teiids, or absence of microchromosomes in (C) geckos and (D) lacertids. Microchromosomes denoted with black bars to the bottom and right of the respective contact map. Annotation of microchromosomal organization denoted via top-right bracket in (A) and (B).

It is difficult to generalize across study systems when using arbitrary numerical cutoffs for what makes a microchromosome in different taxa. We briefly explored this concept in available bird data (chicken, zebra finch, and black swan; Supplemental Figure 1, panels I–K). Specifically, we used two arbitrary cutoffs to group macro/micro chromosomes (1) microchromosomes <30Mb and (2) microchromosomes <10Mb. We can see that chromosomes <10MB possess far more extreme values of gene and GC content than those >10Mb more-or-less meeting the *a priori* expectations of microchromosomal composition. However, using a <30Mb cutoff is more representative of the original karyotypic ‘definition’ of a microchromosome (Boring, 1923). Importantly, when investigating the correlation between chromosome size, GC content, and chromosomal interaction within a single species, the black swan showed a disassociation between chromosome size and (1) higher GC content and (2) chromatin conformation that are both generally associated with microchromosomes (Figure 4a; Supplemental Figure 1, panel K). We see that although a <30Mb cutoff is representative of the karyotypic definition of microchromosome, only chromosomes at a <15Mb cutoff appear to be enriched for the predicted microchromosomal interaction that ‘true’ microchromosomes are expected to possess (Figure 4a; Perry et al. 2021). Thus, it is unclear how to best navigate categorizing chromosomes as macro/micro and the downstream implications on studying the innate properties of these entities.

These inconsistencies bring up a logical conflict as to the nomenclature of microchromosomes. At this point, there are two equally valid ways to ‘define’ a microchromosome, (a) the historical definition of small dot-like chromosomes that are difficult to pair cytogenetically (e.g. Boring, 1923; Hance, 1924) or (b) a grouping of relatively small chromosomes within a genome that possess a distinct nuclear organization (Figure 4; Perry et al. 2021; Waters et al. 2021). It’s important to note that, like in the tegu (Figure 4b), these definitions do not necessarily conflict, however, like in the swan (Figure 4a), they may. By either definition, it is clear that some taxa possess microchromosomes and others do not (Figure 4; Olmo et al. 1990; Perry et al. 2021; Pinto et al. 2022; Srikulnath et al. 2021). Thus, it is important to resolve these conflicts by using specific language that conveys these intricacies. We suggest that rather than attempt to redefine what a microchromosome is *a posteriori*, we qualify the evidence weighted to how we describe microchromosomes. Specifically, at least until we better understand the nuclear function of the observed nuclear organizations of microchromosomes, we can retain the historical definition (1) of microchromosomes and specifically preface those microchromosomes that are isolated in the nucleus as “organized microchromosomes’’. For an example under this framework, the black swan (Figure 4a), all chromosomes <30Mb (10-28) are microchromosomes, but only chromosomes <15Mb would likely be considered “organized microchromosomes’’; however, in the tegu (Figure 4b) all microchromosomes would be considered organized microchromosomes. This type of classification may help clarify communication regarding microchromosomes and any potential functional role these sequestered microchromosomal foci may have across taxa. Further investigations into the evolution of microchromosomes are necessary and likely ongoing, however, to fully understand how microchromosomes evolve the field will need access to additional genomic data from across squamates.

## Conclusion

In conclusion, as prices in genome sequencing continue to fall, squamate genomics will exponentially increase (see also Card et al. 2023). However, keeping up with this progress will not be a trivial task. We show here that the currently available reference genomes, however sparse, are phylogenetically broad enough to make significant contributions to our understanding of genome evolution in vertebrates and additional data will only serve to deepen this understanding. We caution that high-quality genomes without high-quality annotations are limited in their utility to the broader field, but this is an area that needs additional attention from both funding sources, program developers, and empiricists; we see potential for cloud computing as a resource for this work. Current work in sex chromosome and microchromosome evolution (among others) stand to make great strides in coming years as high-quality genomic data become more prevalent in additional taxa. Thus, squamate genomics as a field has blossomed in recent years and this presents a bright outlook for the future of genomics of these often overlooked, yet speciose and charismatic animals.

## Methods

We compiled a near-complete list of all available lepidosaur genome assemblies from GenBank (NCBI), Ensembl (EVI), DNA Zoo (Dudchenko et al. 2017), National Genomics Data Center (CNCB), and individual paper data repositories (e.g. Figshare and GigaDB). We noted the disclosed technologies used to acquire the assembly (from either the database, when available, or the primary article) whether each assembly had an accompanying annotation file available from the download source. We then downloaded each assembly to confirm its existence/availability and calculated basic statistics on each using assembly-stats [v1.0.1] (https://github.com/sanger-pathogens/assembly-stats). We then conducted a literature search to identify the sex determining system of each species (if known), the linkage group (in *Gallus gallus*), the sex-determining region location (if known), and putative sex determining genes. We used this information to assess whether each assembly was considered to be “chromosome-level” or not (in squamates generally, if the scaffold L50 <8 but varies by species) and analyzed this subset using BUSCO [v5.1.2] (Simão et al. 2015) on the gVolante web server [v2.0.0] (Nishimura et al. 2017). Further, for the four genome assemblies of species that were not annotated, which also had an outdated assembly that was annotated, we used Liftoff [v1.6.3] (Shumate and Salzberg, 2021) and uploaded them to an archived repository for public availability and reuse (https://doi.org/10.6084/m9.figshare.20201099). All counts of number of species per clade were collected from Reptile Database (Uetz et al. 2022)

To better understand the genomic composition of reptiles, we used the aforementioned information to best inform which taxa would be the most informative to three downstream analyses. (1) We compiled summary information for representative genomes across amniotes with per chromosome information for number of genes and GC content from NCBI (a lizard, *Podarcis muralis*; two birds, *Gallus gallus* and *Taeniopygia guttata*; a turtle, *Mauremys mutica*; and human, *Homo sapiens*), with a few exceptions that were not directly available through NCBI including gene numbers for other representative squamates: *Shinisaurus crocodilurus* (Liftoff), *Sphaerodactylus townsendi*, and *Naja naja*. In addition, we calculated gene number and GC content for *Alligator sinensis* (Liftoff) and GC content only for *Cygnus atratus*. For each species, we conducted Bayesian and frequentist correlation analyses using JASP (JASP Team, 2022) between each of three variables: GC content, gene number, and chromosome size (the latter two normalized by dividing each by the mean value for each). (2) From a representative number of taxa with chromosome-level reference genomes, we generated a synteny map across squamates, rooted with chicken. We generated corresponding peptide files from each genome using gffread (Pertea and Pertea, 2020) and calculated the synteny map using Genespace (Lovell et al. 2022). (3) We then collated chromatin-contact information from the DNAZoo for a bird (*Cygnus atratus*) and squamate (*Salvator merianae*) and used the recently published contact map from the leopard gecko, *Eublepharis macularius* (Pinto et al. 2023). For *Podarcis muralis*, we had to generate a contact map (that had not been previously published) from the published genome data (PRJNA515813; Andrade et al. 2019). We used Juicer and 3D-DNA to generate the contact map (Dudchenko et al. 2017) and generated images (Figure 4) at a standardized resolution using Juicebox Assembly Tools [v1.11.08] (Durand et al.

2016).

## Data Availability

As indicated above, our cutoff for finding and including additional genomes to the dataset for this paper was July 12th, 2022 and the information used for this study is summarized in Appendix I. However, we have continued aggregating genomes to the summary table beyond this date and have appended them to a live document available here (https://drpintothe2nd.weebly.com/squamates.html). We will continue updating this spreadsheet for the foreseeable future, likely either until genomes become too numerous to keep up with or a better resource is made available. Please feel free to reach out to B.J.P. via email to incorporate additional resources or make corrections to the list.

## Acknowledgements

The authors would like to dedicate this manuscript to Kari Pinto (1960-2022), an avid naturalist, who passed away suddenly during the preparation of this manuscript. This publication was supported by the National Institute of General Medical Sciences of the National Institutes of Health under Award Number R35GM124827 to MAW and NSF IOS 2151318 to TG.

## Supplemental Materials

**Supplemental Figure 1:**
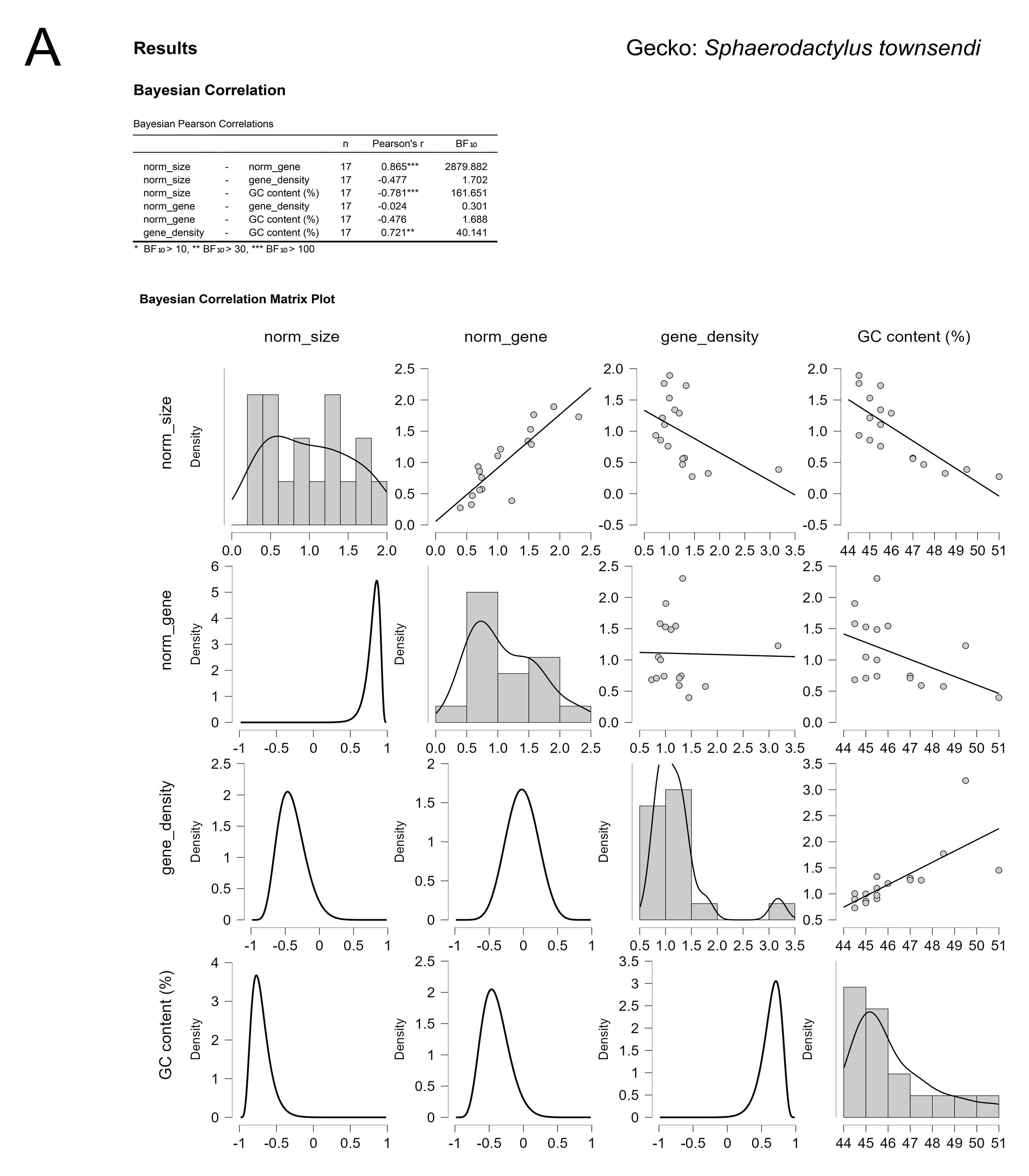
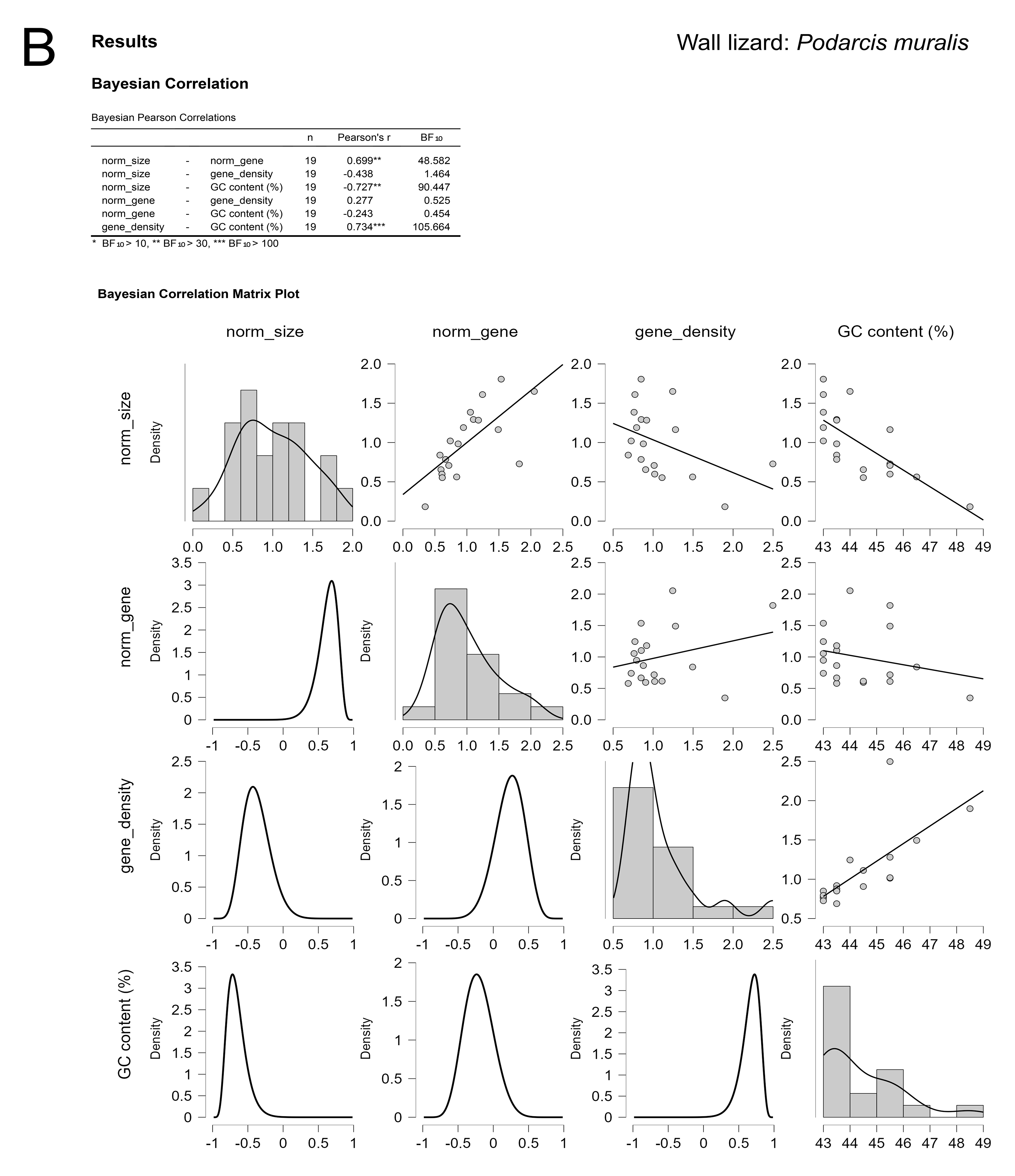
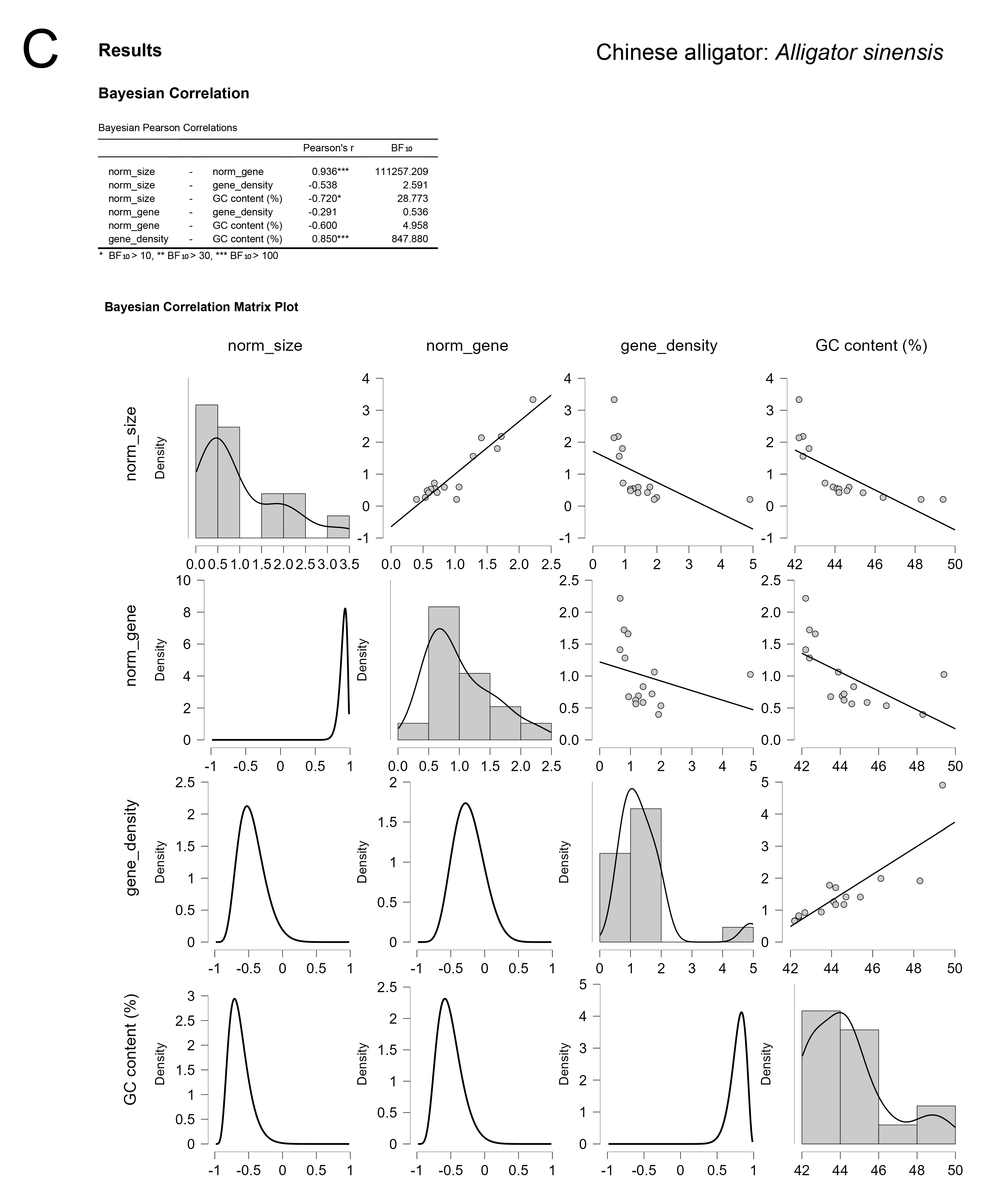
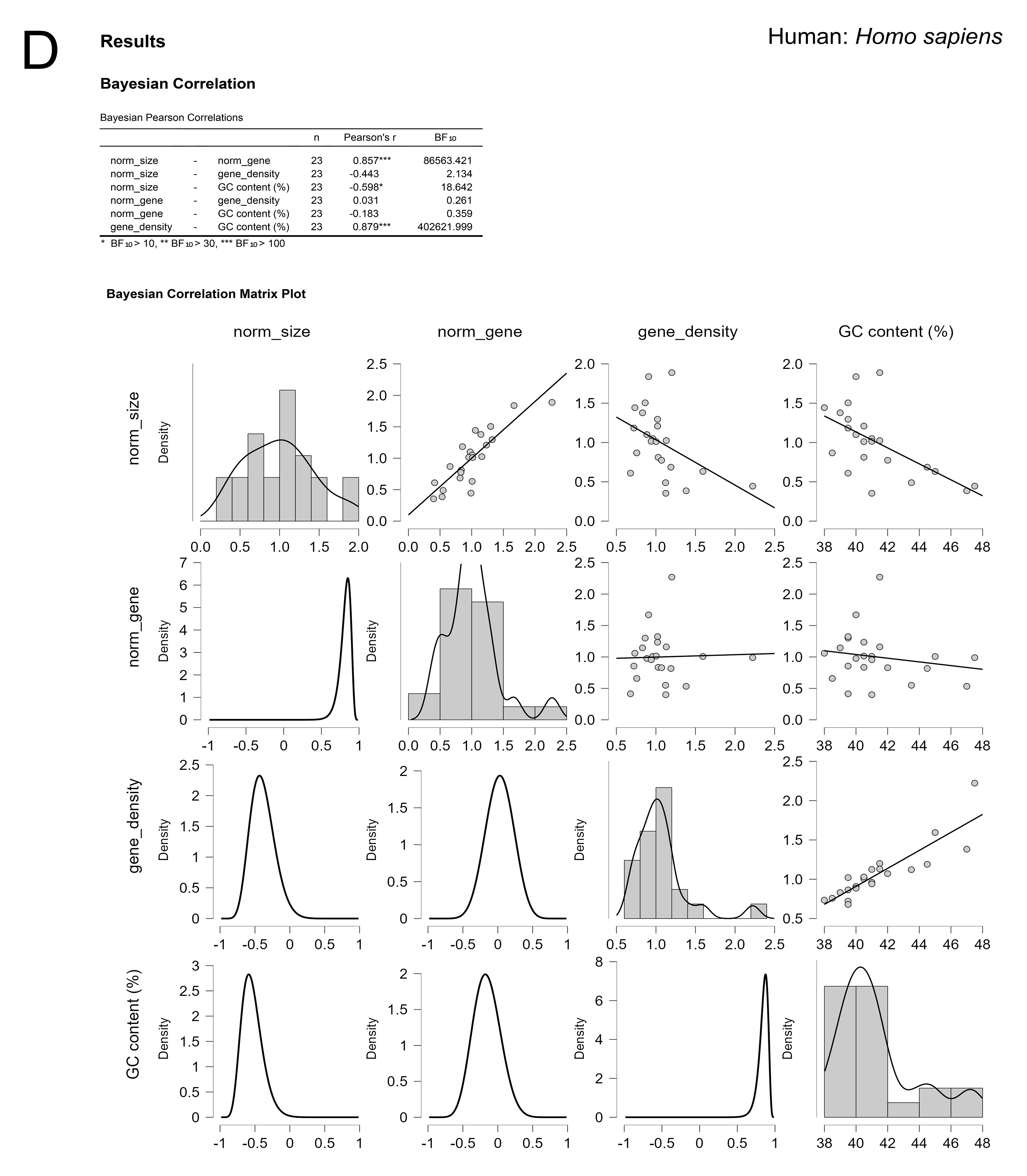
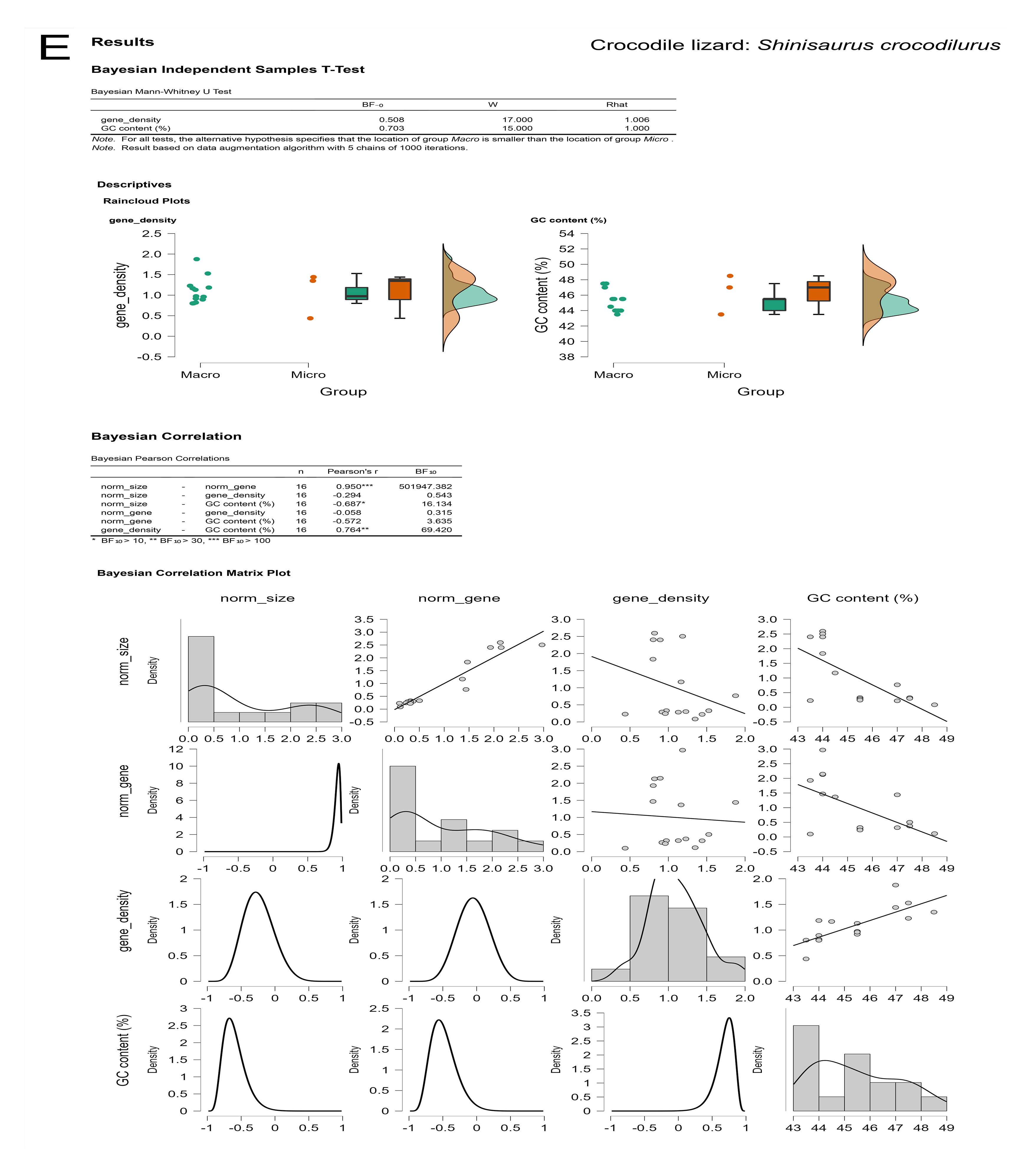
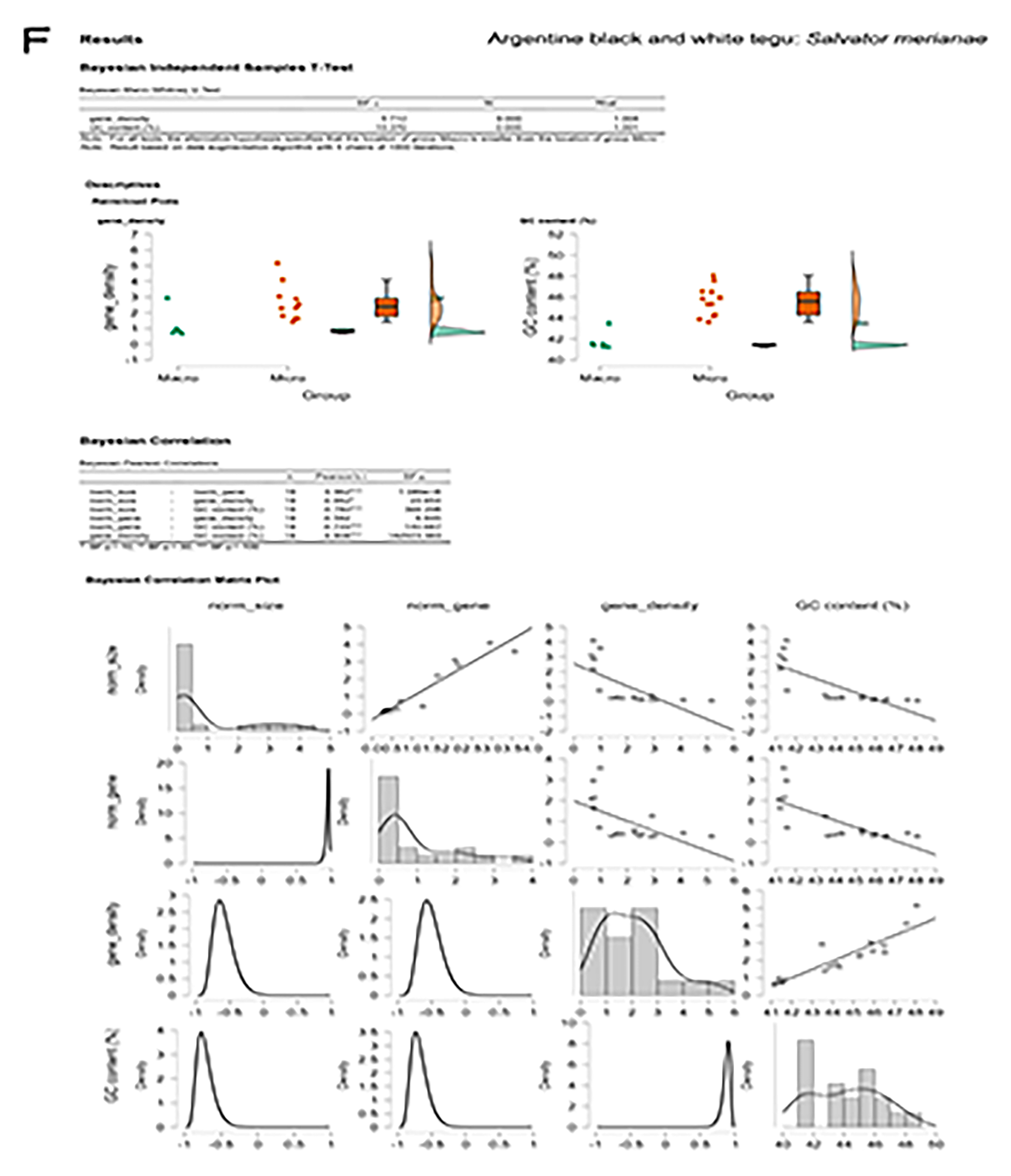
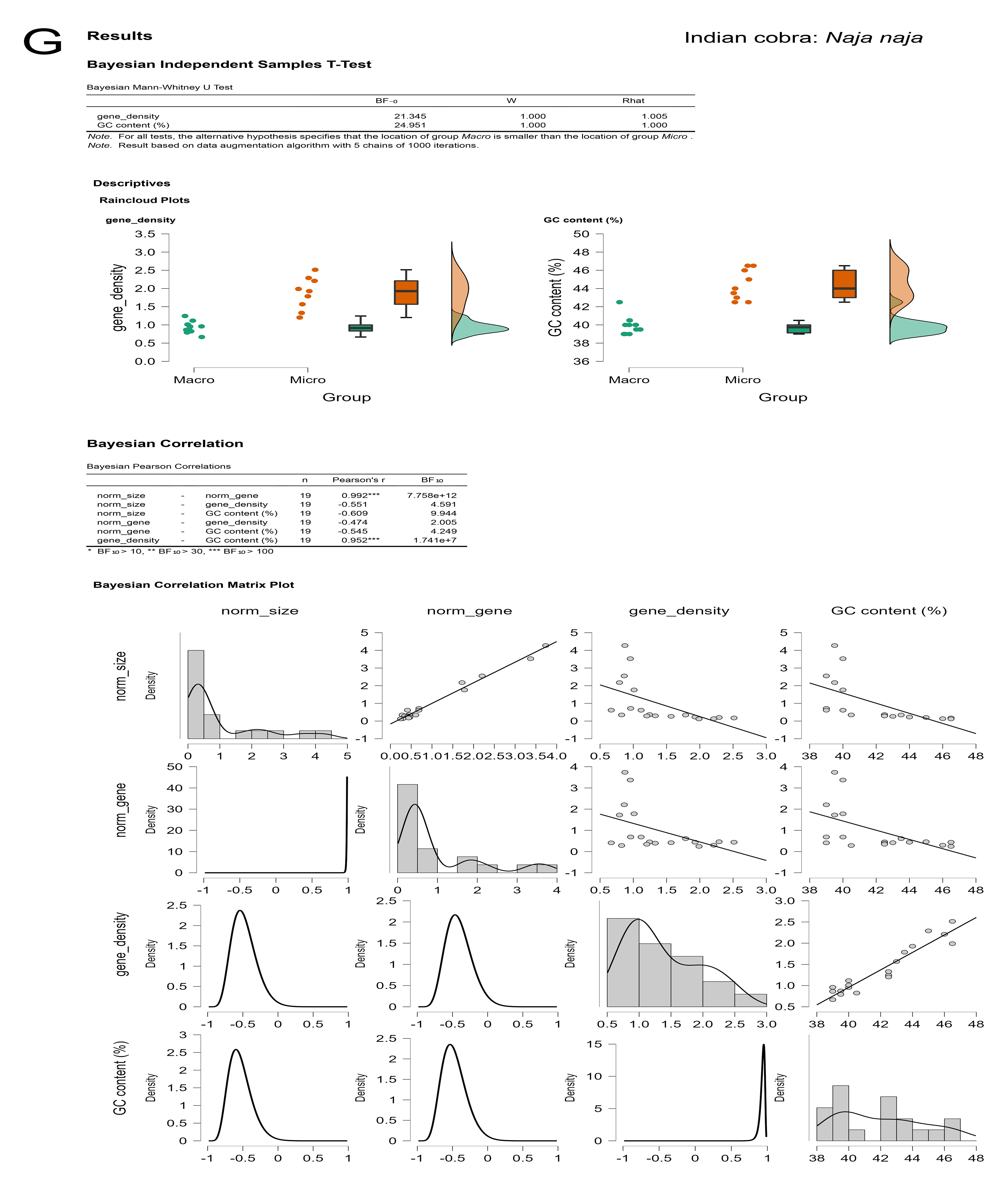
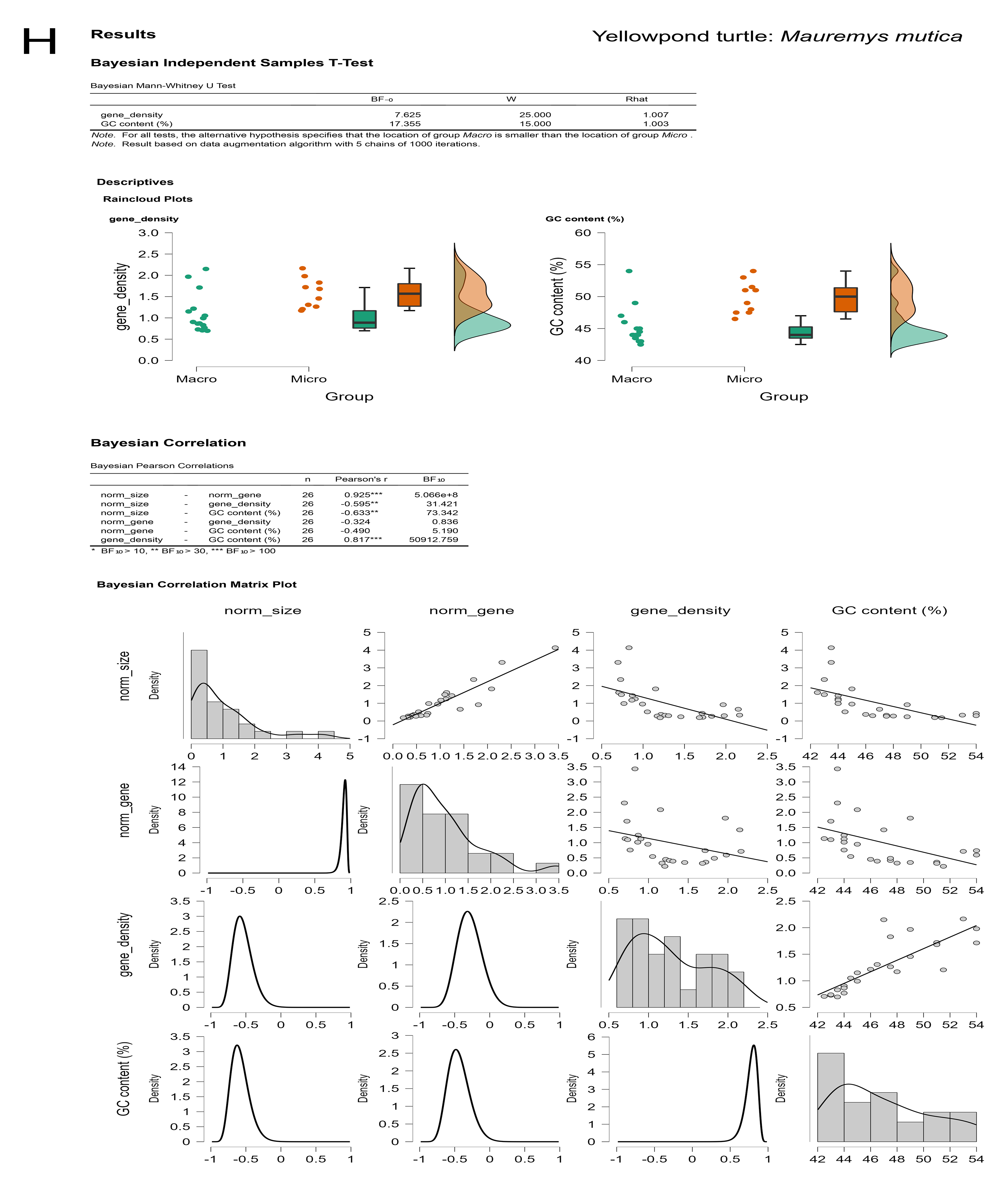
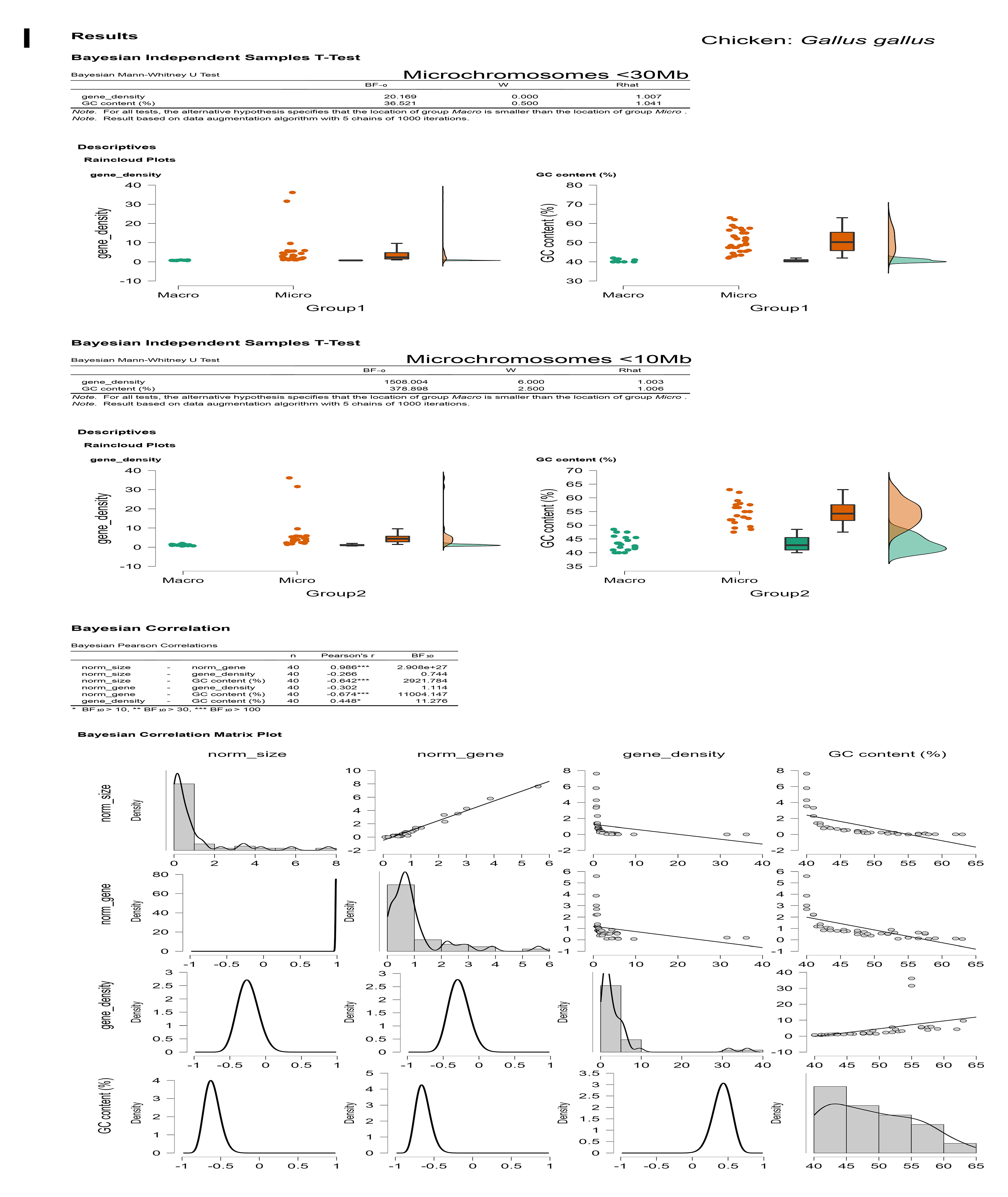
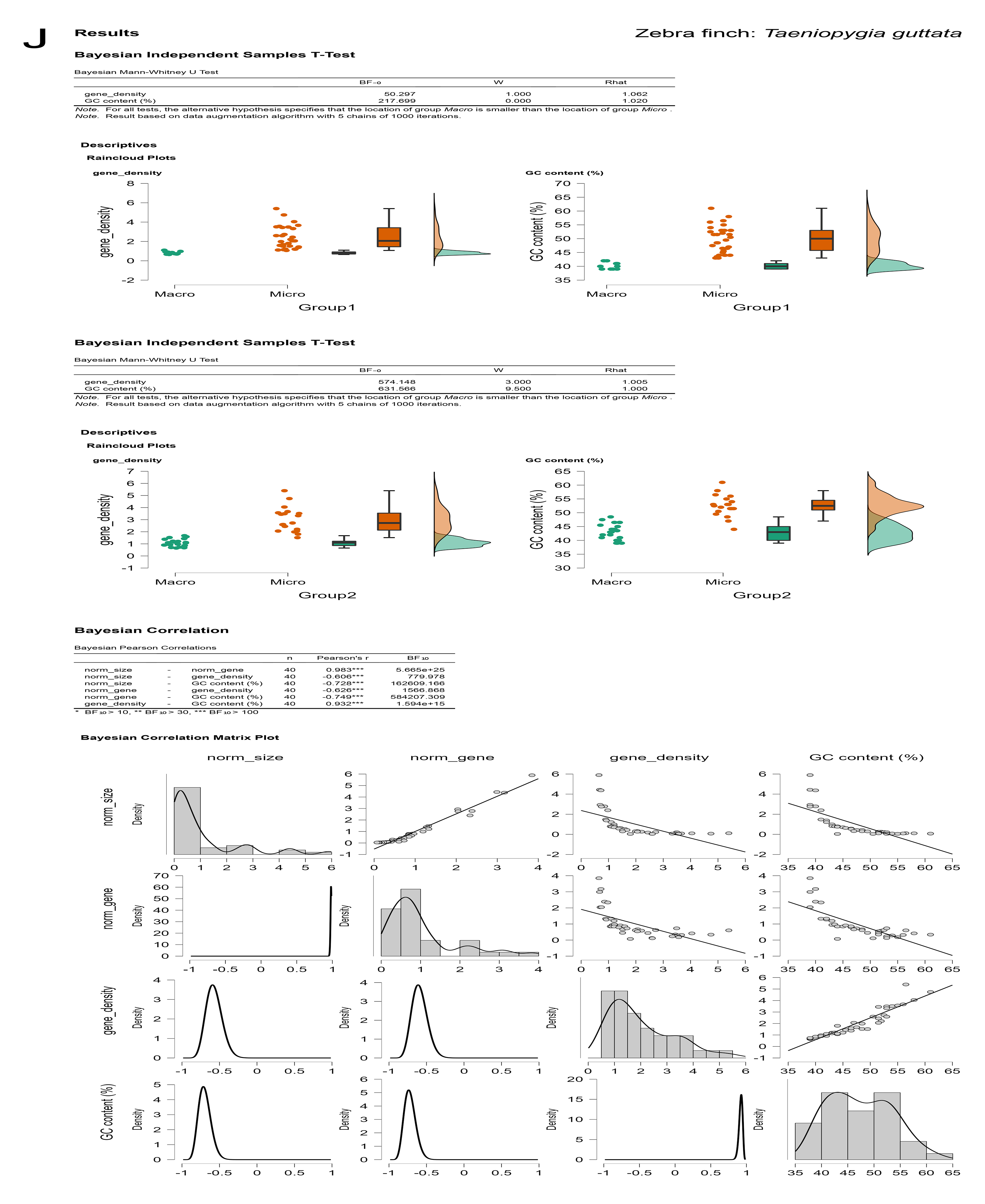
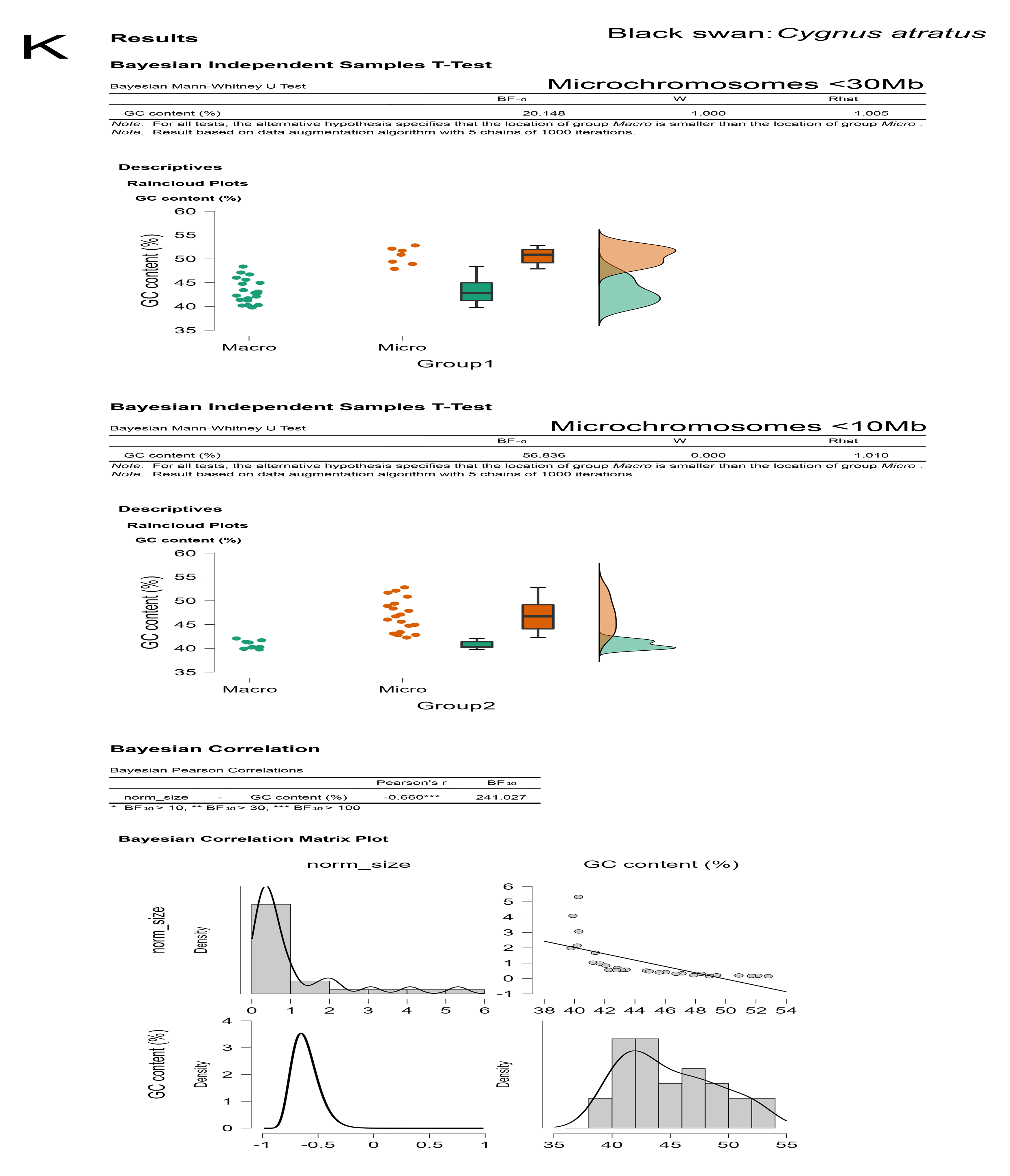
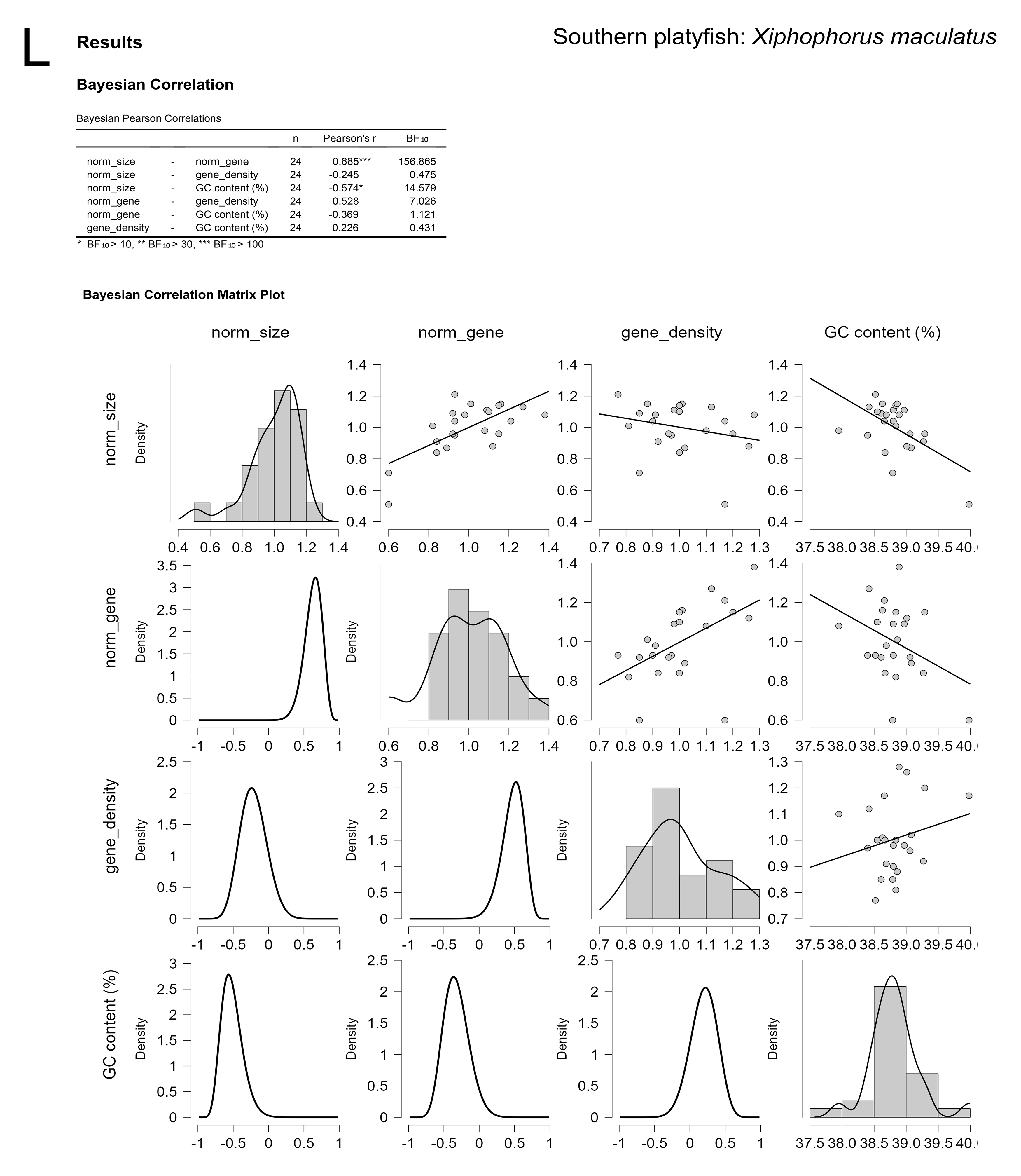
Bayesian analyses of chromosomal make-up across vertebrate taxa. (A) Gecko: *Sphaerodactylus townsendi*, (B) Wall lizard: *Podarcis muralis*, (C) Chinese alligator: *Alligator sinensis*, (D) Human: *Homo sapiens*, (E) Crocodile lizard: *Shinisaurus crocodilurus*, (F) Argentine black and white tegu: *Salvator merianae*, (G) Indian cobra: *Naja naja*, (H) Yellowpond turtle: *Mauremys mutica*, (I) Chicken: *Gallus gallus*, (J) Zebra finch: *Taeniopygia guttata*, (K) Black swan: *Cygnus atratus*, (L) Southern platyfish: *Xiphophorus maculatus*.

**Supplemental Figure 2:**
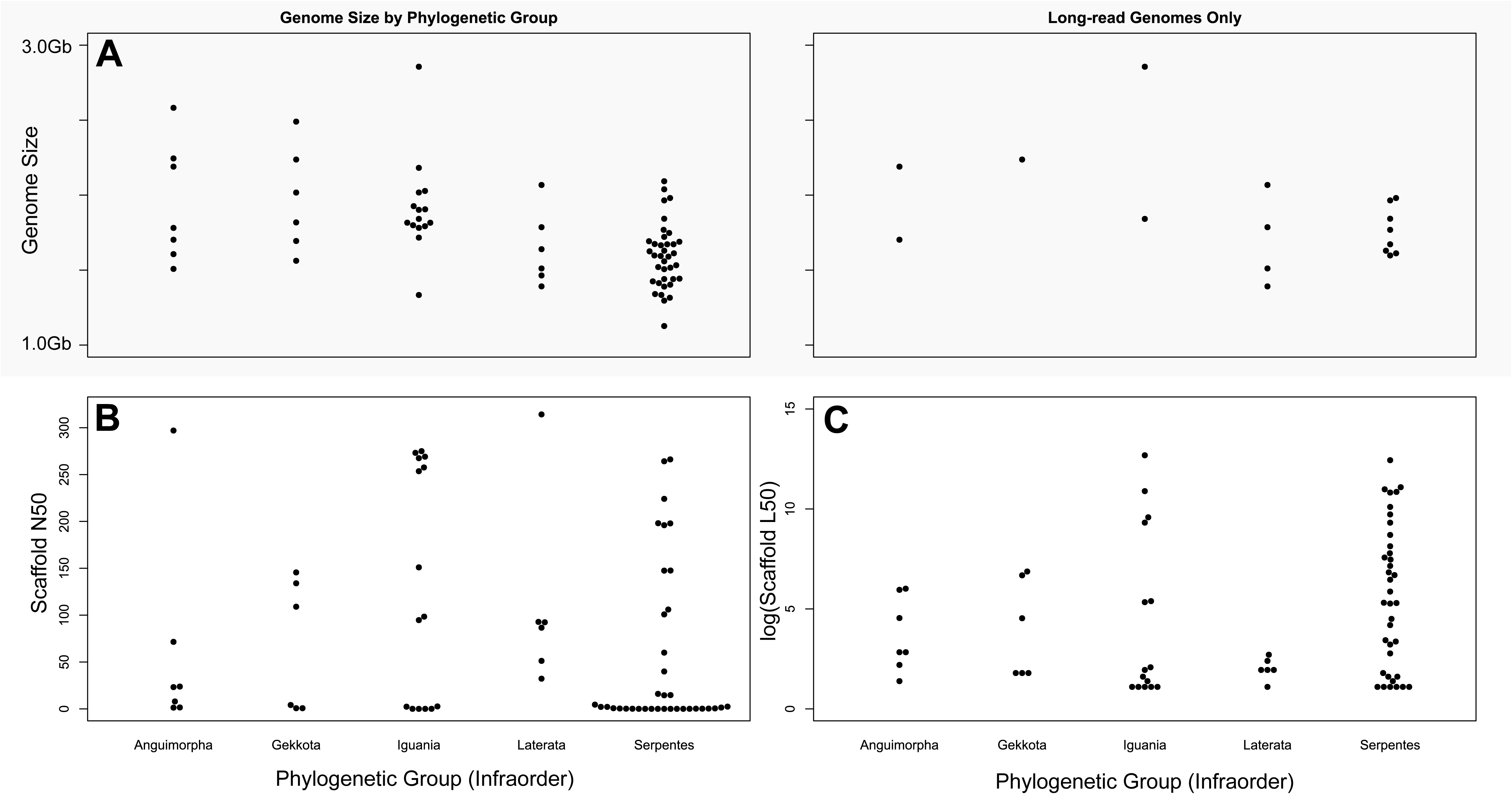
Beeswarm plots splitting genome assemblies by phylogenetic group (Infraorder). (A) Genome size per group for all assemblies (left panel) and long-read only assemblies (right panel). (B) Scaffold N50 and (C) scaffold L50 for all assemblies in each group. Scaffold N50/L50 statistics are capped by physical chromosome sizes within the species in high-quality assemblies, i.e. taxa with macro-/microchromosomes have larger potential N50’s and lower potential L50’s.

**Supplemental Figure 3:**
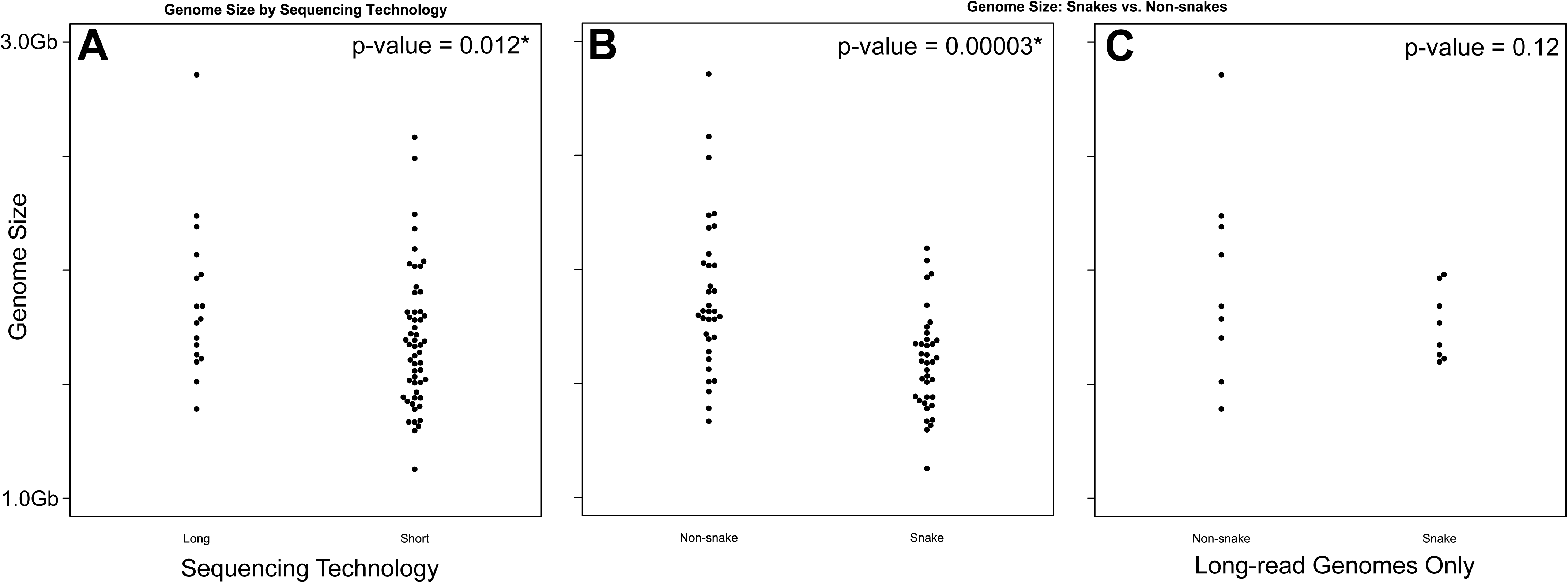
Beeswarm plots splitting genome sizes by technology used to generate the primary assembly (contigs). According to total assembly data (B), snakes appear to have significantly smaller genomes than other squamates. However, (C) when accounting for the extreme bias of short-read assemblies in snakes, this difference disappears. *Long = PacBio and/or ONT, Short = Illumina, Sanger, 454, etc*.

**Supplemental Table 1:**
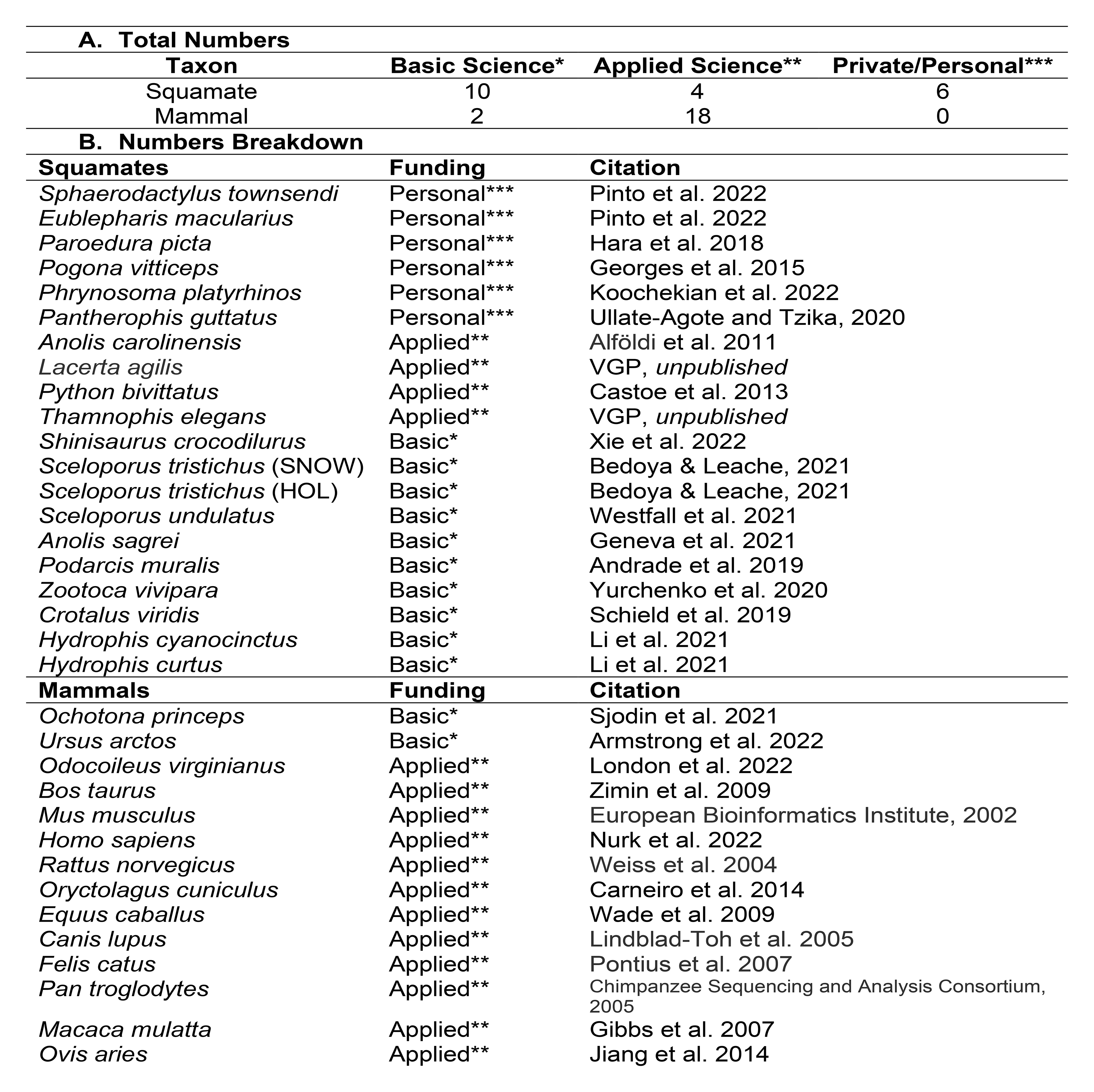
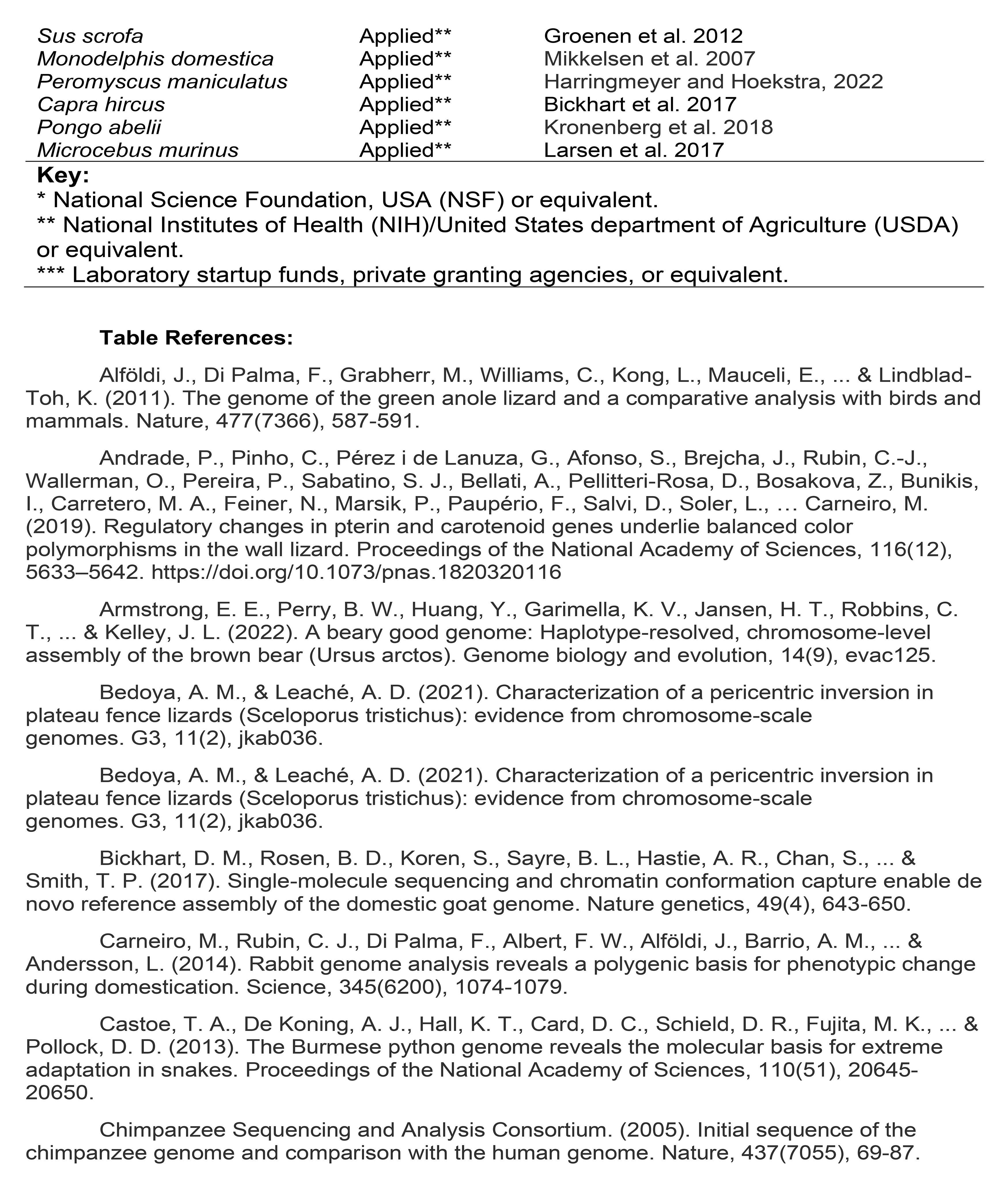
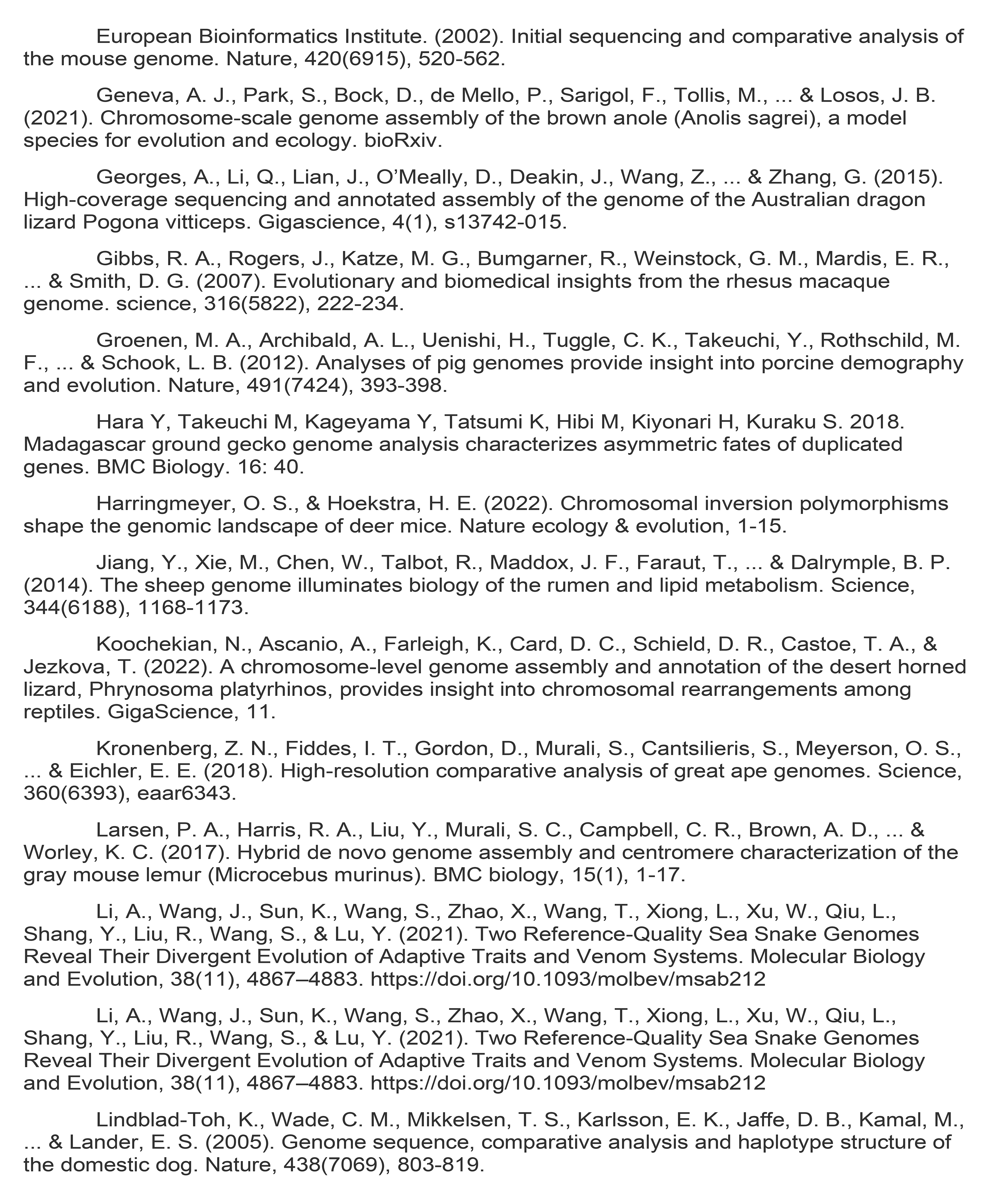
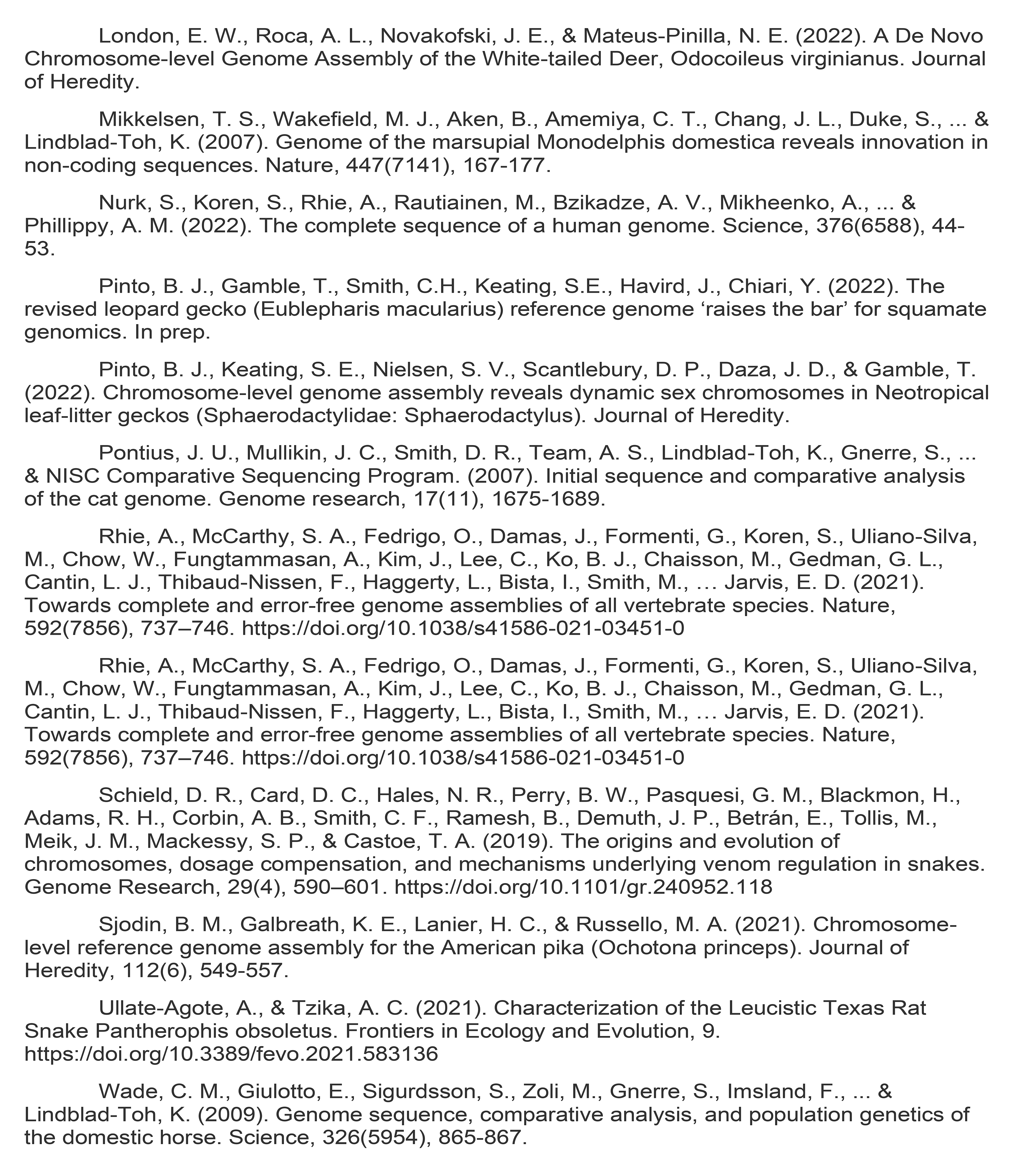
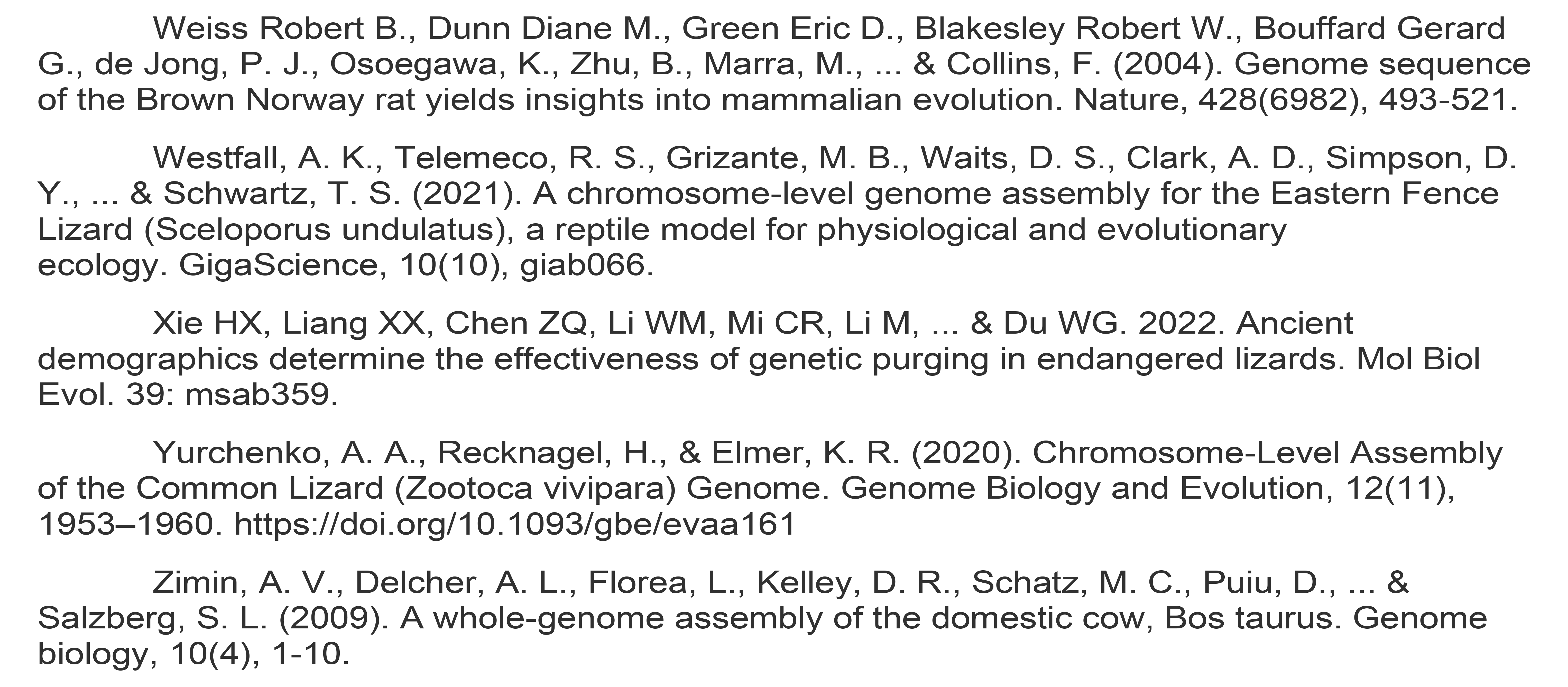
Squamates vs mammal funding source comparison. (A) Most high-quality squamate genomes are generated using soft money from start-up/personal funds or (generally smaller) basic science foundation grants, compared to mammals that receive most of their funding from health and agricultural funding agencies rather than basic/personal funding sources. (B) Breakdown of genomes and references used to generate the funding information shown in A.

## References

1. Abe, T., Kaneko, M., & Kiyonari, H. (2023). A reverse genetic approach in geckos with the CRISPR/Cas9 system by oocyte microinjection. Developmental Biology. 497:26-32.

2. Alföldi, J., Di Palma, F., Grabherr, M., Williams, C., Kong, L., Mauceli, E., … Lindblad-Toh, K. (2011). The genome of the green anole lizard and a comparative analysis with birds and mammals. Nature. 477(7366):587–591.

3. Andrade, P., Pinho, C., Pérez i de Lanuza, G., Afonso, S., Brejcha, J., Rubin, C.- J., Wallerman, O., Pereira, P., Sabatino, S. J., Bellati, A., Pellitteri-Rosa, D., Bosakova, Z., Bunikis, I., Carretero, M. A., Feiner, N., Marsik, P., Paupério, F., Salvi, D., Soler, L., … Carneiro, M. (2019). Regulatory changes in pterin and carotenoid genes underlie balanced color polymorphisms in the wall lizard. PNAS. 116(12):5633–5642.

4. Bachtrog D, Mank JE, Peichel CL, Kirkpatrick M, Otto SP, Ashman TL, Hahn MW, Kitano J, Mayrose I, Ming R, Perrin N, Ross L, Valenzuela N, Vamosi JC. 2014. Sex determination: Why so many ways of doing it? PLoS Biol. 12: e1001899.

5. Blount, Z. D., Lenski, R. E., & Losos, J. B. (2018). Contingency and determinism in evolution: Replaying life’s tape. Science. 362(6415).

6. Bravo, G. A., Schmitt, C. J., Edwards, S. V. (2021). What have we learned from the first 500 avian genomes? Annu Rev Ecol Evol Syst. 52(1):611–639.

7. Bernardi, G., Bernardi, G. (1986). Compositional constraints and genome evolution. Journal of Molecular Evolution. 24(1–2):1–11.

8. Boring, A. M. (1923). Notes by N. M. Stevens on chromosomes of the domestic chicken. Science. 58:73–74.

9. Bull JJ. 1983 Evolution of sex determining mechanisms. Menlo Park, CA: Benjamin Cummings Publishing Company, Inc

10. Bull J, Charnov E. 1985. On irreversible evolution. Evolution 39, 1149–1155.

11. Carey, S. B., Lovell, J. T., Jenkins, J., Leebens-Mack, J., Schmutz, J., Wilson, M. A., & Harkess, A. (2022). Representing sex chromosomes in genome assemblies. Cell Genomics, 100132.

12. Cantarel, B. L., Korf, I., Robb, S. M., Parra, G., Ross, E., Moore, B., … Yandell, M. (2008). MAKER: an easy-to-use annotation pipeline designed for emerging model organism genomes. Genome Research. 18(1):188–196.

13. Card, D. C., Jennings, W. B., & Edwards, S. V. (2023). Genome Evolution and the Future of Phylogenomics of Non-Avian Reptiles. Animals. 13(3):471.

14. Carey, S. B., Lovell, J. T., Jenkins, J., Leebens-Mack, J., Schmutz, J., Wilson, M. A., & Harkess, A. (2022). Representing sex chromosomes in genome assemblies. Cell Genomics. 2(5).

15. Cheng, H., Concepcion, G. T., Feng, X., Zhang, H., Li, H. (2021). Haplotype-resolved de novo assembly using phased assembly graphs with hifiasm. Nature Methods. 18(2):170–175.

16. Cheng, H., Jarvis, E. D., Fedrigo, O., Koepfli, K. P., Urban, L., Gemmell, N. J., Li, H. (2022). Haplotype-resolved assembly of diploid genomes without parental data. Nature Biotechnology. 1-4.

17. Dahn, H. A., Mountcastle, J., Balacco, J., Winkler, S., Bista, I., Schmitt, A. D., Pettersson, O. V., Formenti, G., Oliver, K., Smith, M., Tan, W., Kraus, A., Mac, S., Komoroske, L. M., Lama, T., Crawford, A. J., Murphy, R. W., Brown, S., Scott, A. F., … Fedrigo, O. (2022). Benchmarking ultra-high molecular weight DNA preservation methods for long-read and long-range sequencing. GigaScience. 11.

18. Deakin, J. E., Edwards, M. J., Patel, H., O’Meally, D., Lian, J., Stenhouse, R., Ryan, S., Livernois, A.M., Azad, B., Hollelely, C.E., et al. (2016). Anchoring genome sequence to chromosomes of the central bearded dragon (Pogona vitticeps) enables reconstruction of ancestral squamate macrochromosomes and identifies sequence content of the Z chromosome. BMC Genomics. 17, 1–15.

19. Deakin, J. E., Ezaz, T. (2019). Understanding the Evolution of Reptile Chromosomes through Applications of Combined Cytogenetics and Genomics Approaches. Cytogenetic and Genome Research, 157(1–2), 7–20.

20. Dudchenko O, Batra SS, Omer AD, Nyquist SK, Hoeger M, Durand NC, Shamim MS, et al. (2017). De Novo assembly of the Aedes aegypti genome using Hi-C yields chromosome-length scaffolds. Science. 356:92–95.

21. Durand, N.C., Shamim, M.S., Machol, I., Rao, S.S., Huntley, M.H., Lander, E.S., Aiden, E.L. (2016). Juicer provides a one-click system for analyzing loop-resolution Hi-C experiments. Cell Syst. 3:95–98.

22. Ezaz, T., Sarre, S.D., O’Meally, D., Graves, J.A.M., and Georges, A. (2009). Sex chromosome evolution in lizards: Independent origins and rapid transitions. Cytogenet Genome Res. 127:249–260.

23. Gamble, T. (2010). A review of sex determining mechanisms in geckos (Gekkota: Squamata). Sexual Development. 4(1–2):88–103.

24. Gamble, T., Greenbaum, E., Jackman, T. R., Russell, A. P., & Bauer, A. M. (2012). Repeated origin and loss of adhesive toepads in geckos. PloS One. 7(6): e39429.

25. Gamble, T., Coryell, J., Ezaz, T., Lynch, J., Scantlebury, D. P., Zarkower, D. (2015a). Restriction site-associated DNA sequencing (RAD-seq) reveals an extraordinary number of transitions among gecko sex-determining systems. Molecular Biology and Evolution. 32(5):1296–1309.

26. Gamble, T., Greenbaum, E., Jackman, T. R., & Bauer, A. M. (2015). Into the light: diurnality has evolved multiple times in geckos. Biological Journal of the Linnean Society. 115(4):896–910.

27. Gamble, T., Castoe, T. A., Nielsen, S. V., Banks, J. L., Card, D. C., Schield, D. R., Schuett, G. W., Booth, W. (2017). The discovery of XY sex chromosomes in a *Boa* and *Python*. Current Biology. 27(14):2148–2153.

28. Gamble, T., McKenna, E., Meyer, W., Nielsen, S. V., Pinto, B. J., Scantlebury, D. P., Higham, T. E. (2018). XX/XY sex chromosomes in the South American dwarf gecko (*Gonatodes humeralis*). Journal of Heredity. 109(4):462–468.

29. Gamble, T. 2019. Duplications in corneous beta protein genes and the evolution of gecko adhesion. Integrative and Comparative Biology. 59:193–202.

30. Genome 10K Community of Scientists. (2009). Genome 10K: a proposal to obtain whole-genome sequence for 10 000 vertebrate species. Journal of Heredity, 100(6), 659–674.

31. Gilbert, C., Meik, J. M., Dashevsky, D., Card, D. C., Castoe, T. A., & Schaack, S. (2014). Endogenous hepadnaviruses, bornaviruses and circoviruses in snakes. Proceedings of the Royal Society B: Biological Sciences. 281(1791):20141122.

32. Gu, L., & Walters, J. R. (2017). Evolution of Sex Chromosome Dosage Compensation in Animals: A Beautiful Theory, Undermined by Facts and Bedeviled by Details. Genome Biology and Evolution. 9(9):2461–2476.

33. Fillon, V. (1998). The chicken as a model to study microchromosomes in birds: a review. Genet Sel Evol. 30(3):209–219.

34. Fullerton, S.M., Bernardo Carvalho, A., Clark, A.G. (2001). Local rates of recombination are positively correlated with GC content in the human genome. Mol Biol Evol. 18:1139–1142.

35. Hance, R. T. (1924). The somatic chromosomes of the chick and their possible sex relations. Science. 59:424–425.

36. Hoff, K.J., Lomsadze, A., Borodovsky, M. and Stanke, M. (2019). Whole-Genome Annotation with BRAKER. Methods Mol Biol. 1962:65–95.

37. Hofstra, B., Kulkarni, V. V., Munoz-Najar Galvez, S., He, B., Jurafsky, D., McFarland, D. A. (2020). The diversity–innovation paradox in science. PNAS. 117(17):9284–9291.

38. Hotaling, S., Kelley, J. L., & Frandsen, P. B. (2021). Toward a genome sequence for every animal: Where are we now?. PNAS. 118(52):e2109019118.

39. JASP Team. (2022). JASP (Version 0.16.2) https://jasp-stats.org/

40. Kamiya, T., Kai, W., Tasumi, S., Oka, A., Matsunaga, T., Mizuno, N., Fujita, M., Suetake, H., Suzuki, S., Hosoya, S., Tohari, S., Brenner, S., Miyadai, T., Venkatesh, B., Suzuki, Y., Kikuchi, K. (2012). A Trans-Species Missense SNP in Amhr2 Is Associated with Sex Determination in the Tiger Pufferfish, *Takifugu rubripes* (Fugu). PLoS Genetics. 8(7):e1002798.

41. Karawita, A. C., Cheng, Y., Chew, K. Y., Challgula, A., Kraus, R., Mueller, R. C., Tong, M. Z. W., Hulme, K.D., Beielefeldt-Ohmann, H., Steele, L. E., Wu, M., Sng, J., Noye, E., Bruxner, T. J., Au, G. G., Lowther, S., Blommaert, J., Suh, A., McCauley, A. J., … Short, K. R. (2022). The swan genome and transcriptome: it’s not all black and white. BioRxiv.

42. Keating, S. E., Griffing, A. H., Nielsen, S. V., Scantlebury, D. P., & Gamble, T. (2020). Conserved ZZ/ZW sex chromosomes in Caribbean croaking geckos (*Aristelliger*: Sphaerodactylidae). Journal of Evolutionary Biology, 33(9), 1316–1326.

43. Keating SE. (2022). Evolution of sex chromosomes in geckos (Reptilia: Gekkota). Unpublished dissertation. Marquette University.

44. Keating, S. E., Greenbaum, E., Johnson, J. D., & Gamble, T. (2022).

45. Identification of a cis-sex chromosome transition in banded geckos (Coleonyx, Eublepharidae, Gekkota). Journal of Evolutionary Biology, 35(12), 1675-1682.

46. Kostmann A, Kratochvíl L, Rovatsos M. (2021). Poorly differentiated XX/XY sex chromosomes are widely shared across skink radiation. Proc Roy Soc B. 288:20202139.

47. Kumar S, Stecher G, Suleski M, Hedges SB. 2017. TimeTree: a resource for timelines, timetrees, and divergence times. Mol Biol Evol. 34:1812–1819.

48. Lang, D., Zhang, S., Ren, P., Liang, F., Sun, Z., Meng, G., … Liu, S. (2020). Comparison of the two up-to-date sequencing technologies for genome assembly: HiFi reads of Pacific Biosciences Sequel II system and ultralong reads of Oxford Nanopore. Gigascience. 9(12):giaa123.

49. Li, R., Li, Y., Zheng, H., Luo, R., Zhu, H., Li, Q., Qian, W., Ren, Y., Tian, G., Li, J., Zhou, G., Zhu, X., Wu, H., Qin, J., Jin, X., Li, D., Cao, H., Hu, X., Blanche, H., … Wang, J. (2010). Building the sequence map of the human pan-genome. Nature Biotechnology. 28(1):57–63.

50. Losos J, Braun E, Brown D, Clifton S, Edwards S, Gibson-Brown J, … Warren W. 2005. Proposal to sequence the first reptilian genome: the green anole lizard, *Anolis carolinensis*. NHGRI White Paper.

51. Lovell, J. T., Sreedasyam, A., Schranz, M. E., Wilson, M., Carlson, J. W., Harkess, A., Emms, D., Goodstein, D. M., Schmutz, J. (2022). GENESPACE tracks regions of interest and gene copy number variation across multiple genomes. ELife. 11.

52. Marin, R., Cortez, D., Lamanna, F., Pradeepa, M. M., Leushkin, E., Julien, P., Liechti, A., Halbert, J., Brüning, T., Mössinger, K., Trefzer, T., Conrad, C., Kerver, H. N., Wade, J., Tschopp, P., & Kaessmann, H. (2017). Convergent origination of a Drosophila-like dosage compensation mechanism in a reptile lineage. Genome Research, 27(12), 1974–1987.

53. Meiri, S., Bauer, A. M., Allison, A., Castro-Herrera, F., Chirio, L., Colli, G., … & Roll, U. (2018). Extinct, obscure or imaginary: the lizard species with the smallest ranges. Diversity and Distributions, 24(2), 262–273.

54. Nakamoto, M., Uchino, T., Koshimizu, E., Kuchiishi, Y., Sekiguchi, R., Wang, L., Sudo, R., Endo, M., Guiguen, Y., Schartl, M., Postlethwait, J. H., Sakamoto, T. (2021). A Y-linked anti-Müllerian hormone type-II receptor is the sex-determining gene in ayu, *Plecoglossus altivelis*. PLoS Genetics. 17(8):e1009705.

55. Neemuchwala, S., Johnson, N.A., Pfeiffer, J.M., Lopes-Lima, M., Gomes-dos-Santos, A., Froufe, E., Hillis, D.M., Smith, C.H. 2023. Coevolution with host fishes shapes parasitic life histories in a group of freshwater mussels (Unionidae: Quadrulini).Bulletin of the Society of Systematic Biologists. In press.

56. Newcomer, E. H. (1957). The mitotic chromosomes of the domestic fowl. Journal of Heredity. 48(5):227–234.

57. Nielsen, S. V., Banks, J. L., Diaz, R. E., Trainor, P. A., & Gamble, T. (2018). Dynamic sex chromosomes in Old World chameleons (Squamata: Chamaeleonidae). Journal of Evolutionary Biology, 31(4), 484–490.

58. Nielsen, S. V., Guzmán-Méndez, I. A., Gamble, T., Blumer, M., Pinto, B. J., Kratochvíl, L., & Rovatsos, M. (2019). Escaping the evolutionary trap? Sex chromosome turnover in basilisks and related lizards (Corytophanidae: Squamata). Biology letters, 15(10), 20190498.

59. Nielsen, S. V., Pinto, B. J., Guzmán-Méndez, I. A., & Gamble, T. (2020). First Report of Sex Chromosomes in Night Lizards (Scincoidea: Xantusiidae). Journal of Heredity, 111(3), 307–313.

60. Nishimura O, Hara Y, Kuraku S. (2017). gVolante for standardizing completeness assessment of genome and transcriptome assemblies. Bioinformatics. 33:3635–3637.

61. Nurk, S., Koren, S., Rhie, A., Rautiainen, M., Bzikadze, A. V., Mikheenko, A., Vollger, M. R., Altemose, N., Uralsky, L., Gershman, A., Aganezov, S., Hoyt, S. J., Diekhans, M., Logsdon, G. A., Alonge, M., Antonarakis, S. E., Borchers, M., Bouffard G. G., Brooks, S.Y., … Phillippy, A. M. (2022). The complete sequence of a human genome. Science. 376(6588):44–53.

63. O’Connor RE, Kiazim L, Skinner B, Fonseka G, Joseph S, Jennings R, Larkin DM, Griffin DK. (2019). Patterns of microchromosome organization remain highly conserved throughout avian evolution. Chromosoma. 128:21–29.

64. Ohno, S. (1961). Sex chromosome and microchromosomes of *Gallus domesticus*. Chromosoma. 11:484–498.

65. Ohno S. 1967. Sex chromosomes and sex-linked genes. Berlin, Germany: Springer Verlag.

66. Olmo E. (1986). A. Reptilia. In: John B (ed) Animal cytogenetics 4 Chordata 3. Gebriider Borntraeger, Berlin-Stuttgart.

67. Olmo E, Odierna G, Capriglione T, Cardone A. (1990). DNA and chromosome evolution in lacertid lizards. In: Olmo E, editor Cytogenetics of Amphibians and Reptiles. Berlin, Germany: Birkhauser Verlag. p. 181–204.

68. Olney, K. C., Brotman, S. M., Andrews, J. P., Valverde-Vesling, V. A., & Wilson, M. A. (2020). Reference genome and transcriptome informed by the sex chromosome complement of the sample increase ability to detect sex differences in gene expression from RNA-Seq data. Biology of Sex Differences. 11(1).

69. Palmer B. (2020). Funannotate v1.8.1: Eukaryotic genome annotation pipeline. https://doi.org/10.5281/zenodo.4054262

70. Pan, Q., Herpin, A., & Guiguen, Y. (2022). Inactivation of the Anti-Müllerian Hormone Receptor Type 2 (*amhrII*) Gene in Northern Pike (*Esox lucius*) Results in Male-To-Female Sex Reversal. Sexual Development. 1–6.

71. Pensabene, E., Yurchenko, A., Kratochvíl, L., & Rovatsos, M. (2023). Madagascar Leaf-Tail Geckos (Uroplatus spp.) Share Independently Evolved Differentiated ZZ/ZW Sex Chromosomes. Cells, 12(2), 260.

72. Peona, V., Blom, M. P. K., Xu, L., Burri, R., Sullivan, S., Bunikis, I., Liachko, I., Haryoko, T., Jønsson, K. A., Zhou, Q., Irestedt, M., & Suh, A. (2020). Identifying the causes and consequences of assembly gaps using a multiplatform genome assembly of a bird-of-paradise. Molecular Ecology Resources. 21(1):263–286.

73. Perry, B. W., Schield, D. R., Adams, R. H., Castoe, T. A. (2021). Microchromosomes exhibit distinct features of vertebrate chromosome structure and function with underappreciated ramifications for genome evolution. Molecular Biology and Evolution. 38(3):904–910.

74. Pertea, G., Pertea, M. (2020). GFF Utilities: GffRead and GffCompare. F1000Research. 9:304.

75. Pinto, B. J., Nielsen, S. V., & Gamble, T. (2019a). Transcriptomic data support a nocturnal bottleneck in the ancestor of gecko lizards. Molecular Phylogenetics and Evolution. 141:106639.

76. Pinto, B. J., Card, D. C., Castoe, T. A., Diaz, R. E., Nielsen, S. V., Trainor, P. A., Gamble, T. (2019b). The transcriptome of the veiled chameleon (*Chamaeleo calyptratus*): A resource for studying the evolution and development of vertebrates. Developmental Dynamics. 248(8):702–708.

77. Pinto, B. J., Nielsen, S. V., & Gamble, T. (2019c). Transcriptomic data support a nocturnal bottleneck in the ancestor of gecko lizards. Molecular Phylogenetics and Evolution. 141:106639.

78. Pinto, B. J., Keating, S. E., Nielsen, S. V., Scantlebury, D. P., Daza, J. D., Gamble, T. (2022). Chromosome-level genome assembly reveals dynamic sex chromosomes in neotropical leaf-litter geckos (Sphaerodactylidae: *Sphaerodactylus*). Journal of Heredity. 113(3):272–287.

79. Pinto BJ, Gamble T, Smith CH, Keating SE, Havird J, Chiari Y. (2023a). The revised leopard gecko (*Eublepharis macularius*) reference genome provides insight into the considerations of genome phasing and assembly. Journal of Heredity. In press.

80. Pinto BJ, O’Connor B, Schatz MC, Zarate S, Wilson MA. (2023b). Concerning the eXclusion in human genomics: The choice of sex chromosome representation in the human genome drastically affects number of identified variants. BioRxiv.

81. Pokorná, M., & Kratochvíl, L. (2009). Phylogeny of sex-determining mechanisms in squamate reptiles: are sex chromosomes an evolutionary trap? Zoological Journal of the Linnean Society, 156(1), 168–183.

82. Randhawa, S. S., & Pawar, R. (2021). Fish genomes: Sequencing trends, taxonomy and influence of taxonomy on genome attributes. Journal of Applied Ichthyology. 37(4):553–562.

83. Rasys, A. M., Park, S., Ball, R. E., Alcala, A. J., Lauderdale, J. D., Menke, D. B. (2019). CRISPR-Cas9 gene editing in lizards through microinjection of unfertilized oocytes. Cell Reports. 28(9):2288–2292.

84. Revell LJ, 2012. Phytools: an R package for phylogenetic comparative biology (and other things). Methods Ecol Evol. 3:217–223.

85. Rhoads A, Au KF. (2015). PacBio Sequencing and Its Applications. Genom Proteom Bioinform. 13(5):278–289.

86. Rovatsos, M., Pokorná, M., Altmanová, M., & Kratochvíl, L. (2014). Cretaceous park of sex determination: sex chromosomes are conserved across iguanas. Biology Letters, 10(3), 20131093.

87. Rovatsos, M., Vukić, J., Lymberakis, P., & Kratochvíl, L. (2015). Evolutionary stability of sex chromosomes in snakes. Proceedings of the Royal Society B: Biological Sciences, 282(1821), 20151992.

88. Rovatsos, M., Vukić, J., Mrugała, A., Suwala, G., Lymberakis, P., & Kratochvíl, L. (2019a). Little evidence for switches to environmental sex determination and turnover of sex chromosomes in lacertid lizards. Scientific Reports, 9(1).

89. Rovatsos M, Rehák I, Velenský P, Kratochvíl L. (2019b). Shared Ancient Sex Chromosomes in Varanids, Beaded Lizards, and Alligator Lizards. Mol Biol Evol. 36: 1113–20.

90. Rovatsos, M., Farkačová, K., Altmanová, M., Johnson Pokorná, M., & Kratochvíl, L. (2019c). The rise and fall of differentiated sex chromosomes in geckos. Molecular Ecology, 28(12), 3042–3052.

91. Rovatsos, M., Gamble, T., Nielsen, S. V., Georges, A., Ezaz, T., & Kratochvíl, L. (2021). Do male and female heterogamety really differ in expression regulation? Lack of global dosage balance in pygopodid geckos. Philosophical Transactions of the Royal Society B: Biological Sciences, 376(1833), 20200102.

92. Rovatsos, M., Galoyan, E., Spangenberg, V., Vassilieva, A., & Kratochvíl, L. (2022). XX/XY sex chromosomes in a blind lizard (Dibamidae): Towards understanding the evolution of sex determination in squamates. Journal of Evolutionary Biology, 35(12), 1791–1796.

93. Rupp, S. M., Webster, T. H., Olney, K. C., Hutchins, E. D., Kusumi, K., & Wilson Sayres, M. A. (2017). Evolution of dosage compensation in *Anolis carolinensis*, a reptile with XX/XY chromosomal sex determination. Genome Biology and Evolution. 9(1):231–240.

94. Sackton, T. B., & Clark, N. (2019). Convergent evolution in the genomics era: new insights and directions. Philosophical Transactions of the Royal Society B: Biological Sciences. 374(1777):20190102.

95. Schield, D. R., Card, D. C., Hales, N. R., Perry, B. W., Pasquesi, G. M., Blackmon, H., Adams, R. H., Corbin, A. B., Smith, C. F., Ramesh, B., Demuth, J. P., Betrán, E., Tollis, M., Meik, J. M., Mackessy, S. P., & Castoe, T. A. (2019). The origins and evolution of chromosomes, dosage compensation, and mechanisms underlying venom regulation in snakes. Genome Research, 29(4), 590–601.

96. Shumate A, Salzberg SL. 2021. Liftoff: Accurate mapping of gene annotations. Bioinformatics. 12:1639-1643.

97. Simão FA, Waterhouse RM, Ioannidis P, Kriventseva EV, Zdobnov EM. 2015. BUSCO: assessing genome assembly and annotation completeness with single-copy orthologs. Bioinformatics. 31:3210–3212.

98. Smith, C. H. (2021). A High-Quality Reference Genome for a Parasitic Bivalve with Doubly Uniparental Inheritance (Bivalvia: Unionida). Genome Biology and Evolution. 13(3).

99. Smith, C. H., Pfeiffer, J. M., Johnson, N. A. (2020). Comparative phylogenomics reveal complex evolution of life history strategies in a clade of bivalves with parasitic larvae (Bivalvia: Unionoida: Ambleminae). Cladistics 36(5):505–520.

100. Srikulnath K. 2013. The dynamics of chromosome evolution in reptiles. Thai J Genet. 6:77–79.

101. Srikulnath, K., Ahmad, S. F., Singchat, W., Panthum, T. (2021). Why do some vertebrates have microchromosomes?. Cells. 10(9):2182.

102. Sun, Y., Shang, L., Zhu, Q. H., Fan, L., Guo, L. (2021). Twenty years of plant genome sequencing: Achievements and challenges. Trends in Plant Science.

103. Terao, M., Ogawa, Y., Takada, S., Kajitani, R., Okuno, M., Mochimaru, Y., Matsuoka, K., Itoh, T., Toyoda, A., Kono, T., Jogahara, T., Mizushima, S., & Kuroiwa, A. (2022). Turnover of mammal sex chromosomes in the *Sry*-deficient Amami spiny rat is due to male-specific upregulation of *Sox9*. Proceedings of the National Academy of Sciences, 119(49).

104. Uetz P, Freed P, Aguilar R, Hošek J. (eds.) 2022. The Reptile Database, http://www.reptile-database.org, accessed August 26, 2022.

105. Vicoso, B., Bachtrog, D. (2009). Progress and prospects toward our understanding of the evolution of dosage compensation. Chromosome Research. 17(5).

106. Vicoso, B., Emerson, J. J., Zektser, Y., Mahajan, S., & Bachtrog, D. (2013). Comparative Sex Chromosome Genomics in Snakes: Differentiation, Evolutionary Strata, and Lack of Global Dosage Compensation. PLoS Biology, 11(8), e1001643.

107. Vollger, M. R., Logsdon, G. A., Audano, P. A., Sulovari, A., Porubsky, D., Peluso, P., … & Eichler, E.E. (2020). Improved assembly and variant detection of a haploid human genome using single-molecule, high-fidelity long reads. Annals of Human Genetics. 84(2):125–140.

108. Webster, T. H., Couse, M., Grande, B. M., Karlins, E., Phung, T. N., Richmond, P. A., … Wilson, M.A. (2019). Identifying, understanding, and correcting technical artifacts on the sex chromosomes in next-generation sequencing data. Gigascience. 8(7):giz074.

109. Webster, T.H., Vannan, A., Pinto, B.J., Denbrock, G., Morales, M., Dolby, G.A., Fiddes, I.T., DeNardo, D.F., Wilson, M.A. (2023). Incomplete dosage compensation and lack of dosage balance in the ZZ/ZW Gila monster (*Heloderma suspectum*) revealed by de novo genome assembly. In review.

110. Wetterstrand, K.A. (2021) DNA Sequencing Costs: Data from the NHGRI Genome Sequencing Program (GSP). Available at: www.genome.gov/sequencingcostsdata. Accessed 10-10-2022.

111. Yamashina, M. Y. (1944). Karyotype Studies in Birds I. Comparative morphology of chromosomes in seventeen races of Domestic fowl. Cytologia. 13(3-4):270–296.

